# A recurrent neural network model of prefrontal brain activity during a working memory task

**DOI:** 10.1101/2022.09.02.506349

**Authors:** Emilia P Piwek, Mark G Stokes, Christopher Summerfield

## Abstract

When multiple items are held in short-term memory, cues that retrospectively prioritise one item over another (retro-cues) can facilitate subsequent recall. However, the neural and computational underpinnings of this effect are poorly understood. One recent study recorded neural signals in the macaque lateral prefrontal cortex (LPFC) during a retro-cueing task, contrasting delay-period activity before (pre-cue) and after (post-cue) retrocue onset. They reported that in the *pre-cue* delay, the individual stimuli were maintained in independent subspaces of neural population activity, whereas in the *post-cue* delay, the prioritised items were rotated into a common subspace, potentially allowing a common readout mechanism. To understand how such representational transitions can be learnt through error minimisation, we trained recurrent neural networks (RNNs) with supervision to perform an equivalent cued-recall task. RNNs were presented with two inputs denoting conjunctive colour-location stimuli, followed by a *pre-cue* memory delay, a location retrocue, and a *post-cue* delay. We found that the orthogonal-to-parallel geometry transformation observed in the macaque LPFC emerged naturally in RNNs trained to perform the task. Interestingly, the parallel geometry only developed when the cued information was required to be maintained in short-term memory for several cycles before readout, suggesting that it might confer robustness during maintenance. We extend these findings by analysing the learning dynamics and connectivity patterns of the RNNs, as well as the behaviour of models trained with probabilistic cues, allowing us to make predictions for future studies. Overall, our findings are consistent with recent theoretical accounts which propose that retrocues transform the prioritised memory items into a prospective, action-oriented format.

**Author Summary:** Many real-world scenarios require us to manipulate the contents of memory to guide behaviour. For example, when grocery shopping, initially we might keep all the items from our shopping list in mind (i.e., in our short-term memory). However, once we spot the dairy aisle, we might want to extract, or prioritise, the items from our list that belong to this category. The question of how such prioritisation of memory items is achieved in the brain is a topic of active research. A recent study in monkeys provided evidence that initially, individual memory items are kept from interfering with one another by being encoded by different brain activity patterns. However, following a contextual cue (akin to the dairy aisle sign), the short-term memory representations of the prioritised items are reconfigured into a format that collapses across the irrelevant differences and highlights aspects relevant for action. In this study, we modelled the emergence of these representational changes in an artificial neural network model. Our results help explain how such processes can be learnt from experience and why they might emerge in the biological brain.

## Introduction

Decisions are often based on information that occurred in the recent past. To bridge perception and action across an interval, items need to be maintained in short-term memory [1,2]. In the mammalian neocortex, this memory process may rely on neurons that mutually excite each other in a recurrent circuit [3]. Simulations of this process using recurrent neural networks (RNNs) show how maintenance can rely on self-sustaining attractor states in neural state space. These models can account for empirical phenomena such as persistent delay-period activity and the resistance of memories to external perturbation [4]. When RNNs are trained from random weights with gradient descent (rather than being endowed with coding properties by hand) they can recreate neural motifs that are observed in primates in short-term memory studies. Tools from machine learning can thus offer a normative account of how dynamic memory processes are learned through error minimisation [5,6]. In the current paper, we describe an example in which the activity dynamics in an RNN bear a striking resemblance to those reported in the primate brain.

When humans encode multiple stimuli into short-term memory, retrieval performance is enhanced by cues that reveal which item is more likely to be probed for recall. Surprisingly, this benefit occurs even if cues are presented during the delay period, retrospectively prioritising an item that was encoded in the past [7,8]. At first glance, this seems a puzzling phenomenon because these *retrocues* do not provide additional information about the probed stimulus, so that the benefit appears “out of thin air”. One possibility is that retrocues allow stored memories to be reformatted into a new frame of reference that renders them more robust [9].

One recent study exploited multi-neuronal recordings from macaque lateral prefrontal cortex (LPFC) to understand the neural mechanisms of attention to memory [10]. On each trial, monkeys were shown two coloured squares. During the *pre-cue delay*, they maintained both colours in short term memory, before viewing a retrocue that prioritised one of the two items. Following an additional *post-cue delay*, they reported the cued colour by making a saccade to the corresponding position on a colour wheel (**Fig 1A**). Using multivariate analyses, the authors studied the neural geometry (that is, patterns of similarity in neural states elicited by probed and unprobed items as a function of their colour) during both delay periods. Dissimilarities among neural patterns evoked by each colour scaled with distance around the colour wheel, such that each item occupied a position in neural state space that lay on a ring embedded on a 2D plane. This circular geometry was also seen in human EEG signals for an equivalent task [11].

**Fig 1.**
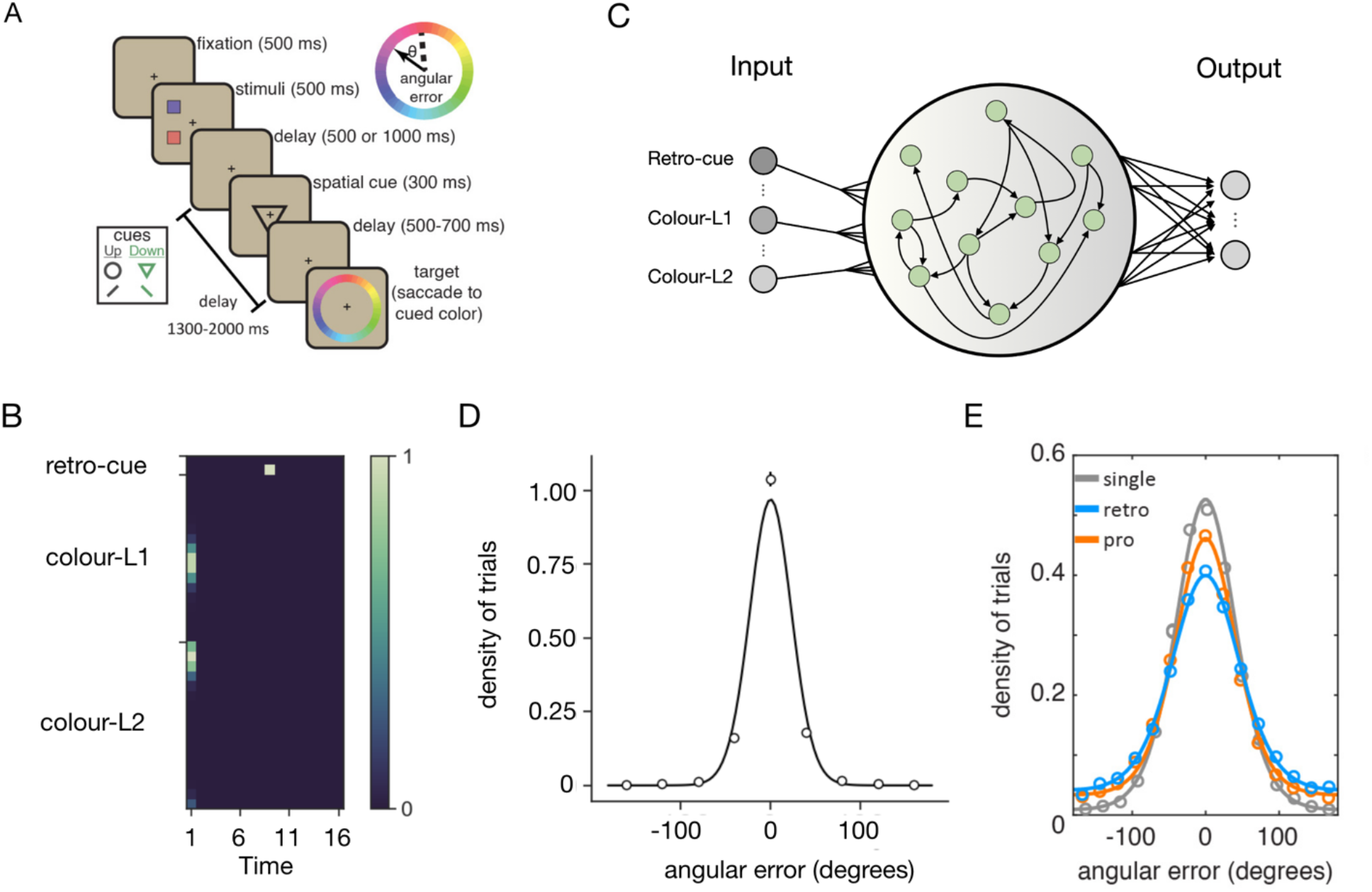
Experimental set-up. **A.** Retro-cueing task structure, taken from [10]. **B.** Input patterns corresponding to an example trial. Colour denotes the activation value of an input unit. Note that inputs near the edges would wrap around because the space is circular **C.** Schematic RNN architecture. Green dots denote recurrent units. Only 3 input units and 2 output are shown (there were in 36 and 17 respectively). **D.** Distribution of errors made by the models at test (circles and vertical bars,*M*±*SEM*) shown with a best fit von Mises (circular normal) density function (line). **E.** Behavioural performance of the monkeys from [10], retro-cueing task shown in blue. Lines correspond to mixture-model fits. Panels A and E are reprinted with permission.

However, the critical finding relates to the orientation of these 2D planes for each delay period. In the pre-cue delay, the two planes lay perpendicular to each other. This is consistent with a growing literature suggesting that the brain minimises interference by encoding information in orthogonal neural subspaces [12–15]. However, in the post-cue delay, the planes rotated so that they lay both in parallel and in phase, forming a common 2D plane whose geometry was isomorphic to the organisation of the colour wheel used for responding [16,17]. This is consistent with the view that retrocues benefit performance by transforming memories into a prospective, action-oriented format, which may render them robust to interference [9].

In the current study, we modelled the emergence of this neural geometry using a recurrent neural network (RNN). We trained randomly initialised RNN models with gradient descent to perform an equivalent retrocue task, predicting that this orthogonal-to-parallel coding geometry would emerge naturally as a consequence of training the network to minimise recall error. This hypothesis was inspired in part by recent findings that neural networks project information into orthogonal subspaces to minimise interference, and parallel subspaces to permit generalisation [12,16–19]. We find that this is indeed the case. We analyse several aspects of the model, including its learning dynamics, its behaviour in the face of probabilistic cues, and its sensitivity to variability in the delay period, allowing us to make predictions for future neural recording studies.

## Results

We report results from 4 simulations (denoted experiments). In Experiment 1, we explore the ability of recurrent neural networks to recreate the data described in [10]. In Experiment 2, we adapt the retrocue paradigm to a new case with variable delay periods and explore how this affects the neural geometry in RNNs. In Experiment 3, we use tools from cognitive psychology to characterize the behaviour of neural networks in a variant of the paradigm in which cues have variable validity, as well as examine relationship between neural geometry and behaviour in the RNNs. Finally, in Experiment 4, we study why and when the geometry emerges.

### Experiment 1

We trained recurrent neural network models (RNNs, N = 30 models with different initialisation and trial order) to perform a retro-cueing task comparable to that previously described in [10] (see **Fig 1A** and **B**). Each model was a vanilla RNN with a single recurrent layer, incorporating rectified linear unit (ReLU) nonlinearities, and a single decoding layer (see **Fig 1C** for a schematic of the network architecture). Each trial was composed of four successive inputs. Firstly, networks received two concatenated circular normal inputs (corresponding to distributed representations of the colour stimuli at two different locations, which we denote *colour-L1, colour-L2*), followed by seven cycles of delay (*pre-cue delay*). This was followed by a one-hot input (the *retro-cue*) denoting which of the two stimulus locations was relevant on the current trial and then finally, a second memory delay (*post-cue delay*). An example trial is shown in **Fig 1B**.

The networks were trained using stochastic gradient descent to minimise the mean squared error (MSE) between their output and the stimulus colour shown at the cued location. Thus, we do not engineer the representations into the networks but allow them to form autonomously during optimisation. To bound performance we added zero-mean Gaussian noise to the hidden activations of the models and trained until a plateau in the loss curve had been detected (see *Methods*). After approximately 220 ± 105 epochs the networks converged to a mean absolute angular error of 16.40 ± 0.94° (*M*±*SEM*). This was somewhat smaller than the value reported for monkeys (51.8°). This discrepancy could be seen to reflect other sources of noise (e.g., motor error during saccadic responding) that are present in the animals but not in our models but is otherwise not consequential for our findings (increasing the noise added to network units precluded convergence).

In Experiment 1, we first explored the representational geometry that related both cued and uncued items in the recurrent layer of the RNN, and compared it to that reported in macaque LPFC [10]. Noting a correspondence between the two, we then studied how this geometry emerged in the RNN over the course of learning, what connectivity patterns gave rise to it, and how it related to network behaviour, allowing us to form predictions for future studies.

#### Representation of cued items

To study the geometry of the cued memory representations formed in the pre-cue and post-cue delay periods, we focused on the “selection” task described in [10]. The geometry was computed by applying dimensionality reduction to the features (neural activity over units or neurons) × conditions (each item binned by colour and cue) matrix and plotting the reduced reconstruction in 3D. We begin by alerting the reader to two frames of reference for this analysis. Firstly, we can ask about the relationship between *colour-L1* and *colour-L2* items when they are cued (*Cued Geometry* below) or uncued (*Uncued Geometry* below). Alternatively, we can ask about the relationship between the two items when one of them is cued and the other is not (*Cued/Uncued Geometry*).

We focused first on the *Cued Geometry*. In monkey LPFC, in both the pre- and post-cue periods, the neural representations arranged themselves in two rings, representing similarity in the circular colour space for each of the two respective locations (L1 and L2; [10]. During the pre-cue delay, the angle between the two planes that lay parallel to the shank of each ring was close to 90°. However, during the post-cue delay the two rings became aligned and phase matched, i.e., corresponding points in colour space were aligned on the two rings. A visualisation of this orthogonal-to-parallel transformation from the paper is shown in **Fig 2A**.

**Fig 2.**
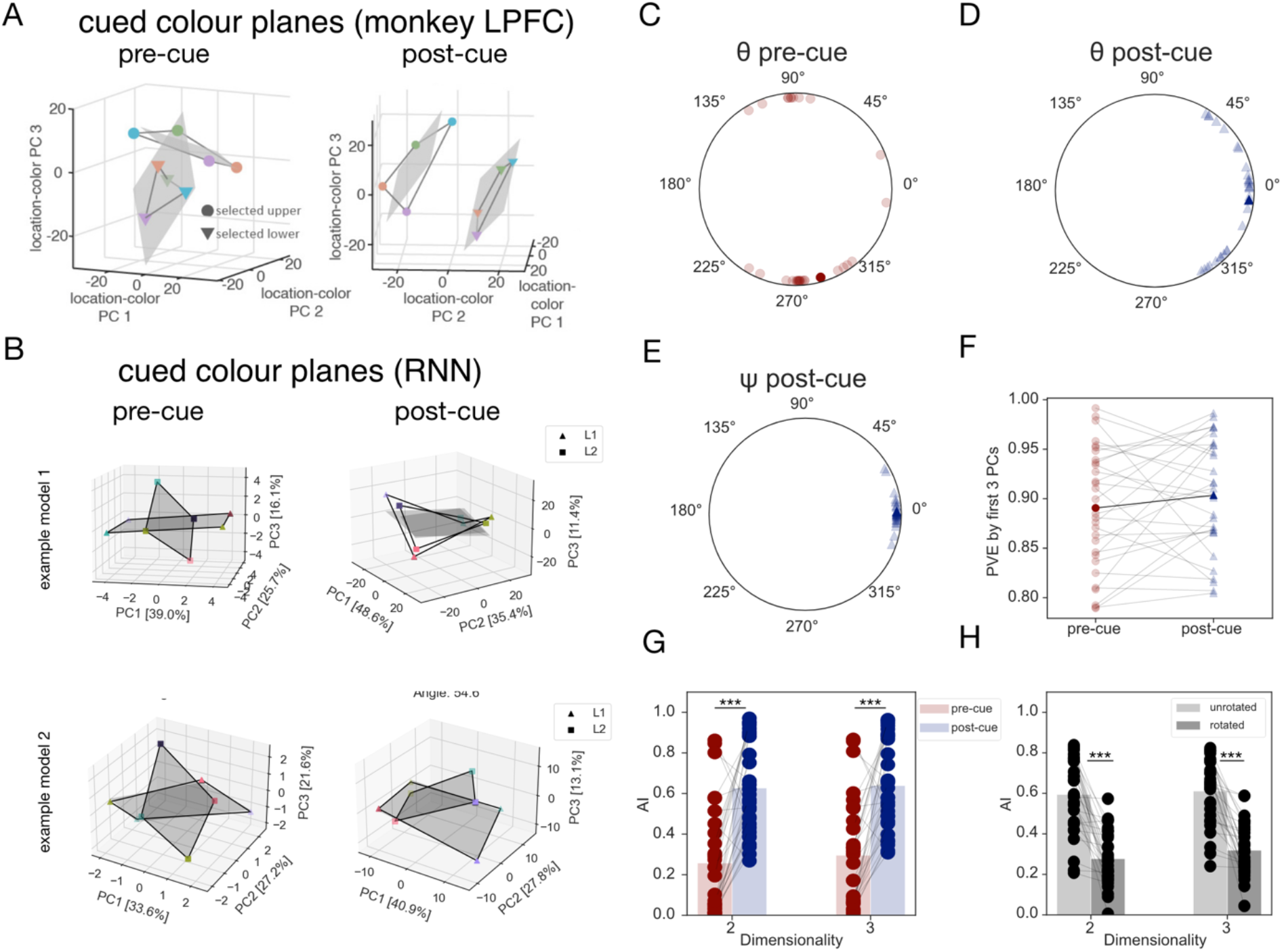
The geometry of cued mnemonic representations learnt by the RNNs. **A.** Visualisation of the pre- (*left*) and post-cue (*right*) geometry of selected (i.e., cued) items reported in [10]. Population responses from LPFC binned into 4 colour categories (denoted by marker colour) for each stimulus location (upper, lower – denoted by marker shape). Data plotted in reduced dimensionality space, defined by the first 3 principal components (PCs) of the 8 location-colour pairs. Planes of best fit for each location shown as grey quadrilaterals. **B.** Analogous visualisation of the hidden activity patterns from two example RNN models for the pre-cue (*left*) and post-cue (*right*) delay. Two locations (L1, L2) correspond to triangles and squares, respectively. The percentage of total variance explained by each PC axis shown in square brackets. Chosen models (in this and subsequent figures) correspond to the ones with geometries qualitatively most and least similar to the group-level average (in this case, the average post-cue geometry). **C.-D.** Between-plane angles *θ* for the pre- (red dots) and post-cue delay (navy triangles) timepoints, respectively. Lighter and darker colours correspond to the values for individual models and grand averages, respectively. **E.** Phase alignment angles ψ between the cued planes in the post-cue delay. **F.** Proportion of total variance explained (PVE) by the first 3 PCs for the PCA models fit to activation patterns from individual networks. Values for individual models shown in lighter, and grand averages in darker colours. **G.** Subspace alignment index (AI) between location-specific planes during the pre- (red) and post-cue (navy) delay intervals. Individual models shown as points, bars correspond to the grand averages. AI reported for 2- and 3-dimensional subspaces **H.** AI for the *unrotated* (light grey) and *rotated* (dark grey) subspaces – for description, refer to the main text. Statistically significant contrasts denoted by asterisks (*: p < .05, **: p < .01, ***: p < .001). Panel A is reprinted with permission.

To study the memory representations created by the trained RNNs, we followed the same approach. We extracted hidden unit activity patterns corresponding to each trial type (defined by the unique colour-L1, colour-L2 and retro-cue combination) at the final timepoint of each delay period. For visualisation shown in **Fig 2B**, we binned the activation patterns corresponding to different stimulus colours into 4 equally spaced colour categories and collapsed the data across the uncued stimuli, resulting in a 200 × 8 (unit activity by conditions) matrix. Subsequently, we performed principal components analysis (PCA) to project the data into a 3-dimensional space.

As reported in [10], we observed that the input representations were structured by their similarity, with the “colour” representations for each stimulus location largely confined to 2-dimensional subspaces. Critically however, visual inspection suggested that the orientation of the location-specific planes in the 3-dimensional space differed between the two trial periods in a very similar way to that seen in monkey LPFC. During the pre-cue delay, the location-specific planes were orthogonal to one another, whereas during the post-cue delay, the two planes had a roughly parallel configuration. Moreover, the corresponding colour category representations were also phase-aligned, potentially permitting a common readout mechanism (see **Fig 2B**).

To quantify the described geometry, we reduced the data to three dimensions for each network, estimated best-fitting planes for the datapoints corresponding to each cued location and calculated the angle *θ* between them in this low-rank space. Across all trained models, *θ* was clustered at the +90° and −90° marks in the pre-cue delay (**Fig 2C**). To test for the statistical significance of this effect, we rectified the data and performed a v-test with a hypothesised angular mean of +90° (see *Methods* for more details). This returned a significant result (*v(29)* = 22.18, *p* < .001), confirming that the pre-cue planes were indeed orthogonal. In contrast, in the post-cue delay, *θ* was significantly clustered around 0° (circular mean = −5.66°; v-test results: *v(29)* = 24.09,*p* < .001, see also **Fig 2D**). The mean absolute difference between pre- and post-cue *θ* was also significantly different from 0° (one-sample test for mean angle, *M* = 93.2°, 95% *CI* = 69.66°, 116.74°).

In ref [10], the authors report that the planes were phase aligned during the post-cue period. To quantify the phase alignment between the L1 and L2 datapoints in the RNN, we projected them onto their respective planes before rotating the planes to be coplanar. The phase alignment angle ψ was then obtained from a rotation matrix that most closely mapped L1 datapoints onto L2 datapoints. Importantly, ψ was only defined for geometries that could not be classified as either mirror reflections or bowtie shapes. Given that the pre-cue planes were significantly orthogonal, we only report ψ for the post-cue delay, where it was defined for all models and significantly clustered around 0° (circular mean = 1.67°, v-test results: *z*(29) = 29.57, p < .001, *N* = 30). Thus, our results indicate that in the RNN, like in the monkeys, the post-cue planes were both parallel and phase-aligned.

We confirmed the above results with an analogous analysis using the subspace alignment index (AI) introduced in [20] (see also *Methods*). This method projects the data from condition *i* into the subspace of condition *j* and calculates the amount of variance of *i* explained by the subspace of *j*. The AI metric takes values between 0 and 1, which correspond respectively to fully orthogonal and fully aligned subspaces. We calculated the AI for 2 and 3 dimensional subspaces and found that in both cases, the mean pre-cue AI was low (AI =0.26 and 0.3, for 2 and 3 dimensions, respectively), indicating nearly orthogonal subspaces for the two locations. In contrast, the mean post-cue AI was significantly higher (more parallel) in both target dimensionalities (2D: AI = 0.63, paired t-test: *t*(29) = 6.13,*p* < .001, one-tailed; 3D: AI = 0.64; *t*(29) = 5.98, *p* < .001, one-tailed), indicating a significant increase in subspace alignment across the two delays (see also **Fig 2G**).

Lastly, we investigated how a given colour ring was oriented with respect to itself before and after the presentation of the retro-cue. This allowed us to study the nature of the transformation that the network had learned to perform. To parallelise the subspaces, the networks could either learn to rotate both pre-cue subspaces, or to rotate only one, keeping the other static. To establish which solution was utilised by the models, we performed the AI analysis in 2-fold cross-validation. This was necessary because the frame of reference of location (e.g., left vs. right side) is arbitrary, and so if different networks learned to rotate left or right stimuli, averaging the data would obscure the results. Thus, we used half of the data to identify the *unrotated* and *rotated* locations (based on their AI values), and the other half to confirm whether there was a significant difference between them. We observed that the mean AI for the *unrotated* location (AI = 0.60 and 0.61 for 2 and 3D, respectively) was significantly higher than for the *rotated* location (2D: AI = 0.28, Wilcoxon signed-rank test *W*(29) = 462, *p* < .001; 3D: AI = 0.32; *W*(29) = 465, *p* < .001; see **Fig 2F**) and comparable to the AI for the post-cue subspaces (**Fig 2G**). This implies that by convergence, the networks had learnt to rotate only one of the pre-cue subspaces, keeping the other in its pre-cue orientation. These findings diverge from the observations made in monkey LPFC, where the different task periods were associated with independent subspaces.

#### Representation of uncued items - *Uncued Geometry*

The RNN thus seems to capture the cued stimulus representations seen in macaque prefrontal cortex. Next, we turned to investigate the *Uncued Geometry*, that is the relationship between items *colour-L1* and *colour-L2* when they were not signalled by the retrocue. First, we asked whether the RNNs retained any information at all about the uncued item in the post-cue period. In monkey LPFC, there was significant colour information about both memory items (quantified as the circular entropy of each neuron’s response to colour) in all trial periods [10]. This was the case even though the monkeys were never probed on the uncued memory item and could have, in theory, discarded it after the cue presentation. We found the same to be true in the RNNs. For all networks, linear decoders trained to perform pair-wise colour discriminations of the uncued items in the post-cue delay showed significantly above-chance mean performance when tested on held out data (*M* = 62.46% decoding accuracy, one-sample t-test, *t*(29) = 15.47,*p* < .001).

We then set to find out if the uncued memory items also underwent a transformation from a feature-based representation to a more response-oriented format. In monkey LPFC, the uncued subspaces became slightly, but non-significantly, more parallel in the post-cue delay. Moreover, the two subspaces, unlike their cued counterparts, were not phase-aligned [10] (**Fig 3A**). To visualise and quantify the geometry of these representations across our models, we followed the same steps as described above (in the section subtitled *representation of cued items*), except the data were averaged across uncued rather than cued locations (the configuration in the post-cue delay shown in **Fig 3B**). For each delay period, we calculated and compared the two location subspaces using the AI (**Fig 3D**). The mean AI in the post-cue delay was marginally higher than in the pre-cue delay (2D: pre-cue AI = 0.26, post-cue AI = 0.32, paired t-test: *t*(29) = 1.36, *p* = .092, one-tailed; 3D: pre-cue AI = 0.30, post-cue AI = 0.37, paired t-test: *t*(29) = 1.77, *p* = .043, one-tailed). It was also, however, significantly lower than the post-cue index for the cued subspaces (paired t-test: 2D: *t*(29) = −6.74, *p* < .001; 3D: *t*(29) = −5.84,*p*< .001). A complementary analysis of the angles formed by the two planes in the post-cue delay revealed that their orientation did not show any consistent trend across trained models (see **Fig 3C**, *left*). This was confirmed by the results of a Rayleigh test which failed to reject the null hypothesis of a uniform distribution of plane angles around the unit circle (*z*(29) = 0.06, *p* = .941). Similarly, there was no significant phase alignment between the uncued planes (v-test results: v(15) = 1.81, *p* = .261, *N* = 16), and nearly 50% of the models developed geometries where the two planes were either mirror reflections of one another, or one of the geometries was twisted into a bowtie shape. Taken all together, these findings show that the two uncued subspaces are generally not transformed into a common frame of reference post-cue and their precise arrangement is variable between the trained models. Whilst one might be tempted to propose that the *Uncued geometry* is irrelevant for the choice, we shall see below that (perhaps surprisingly) it is reliably predictive of behaviour (see Experiment 3 below).

**Fig 3.**
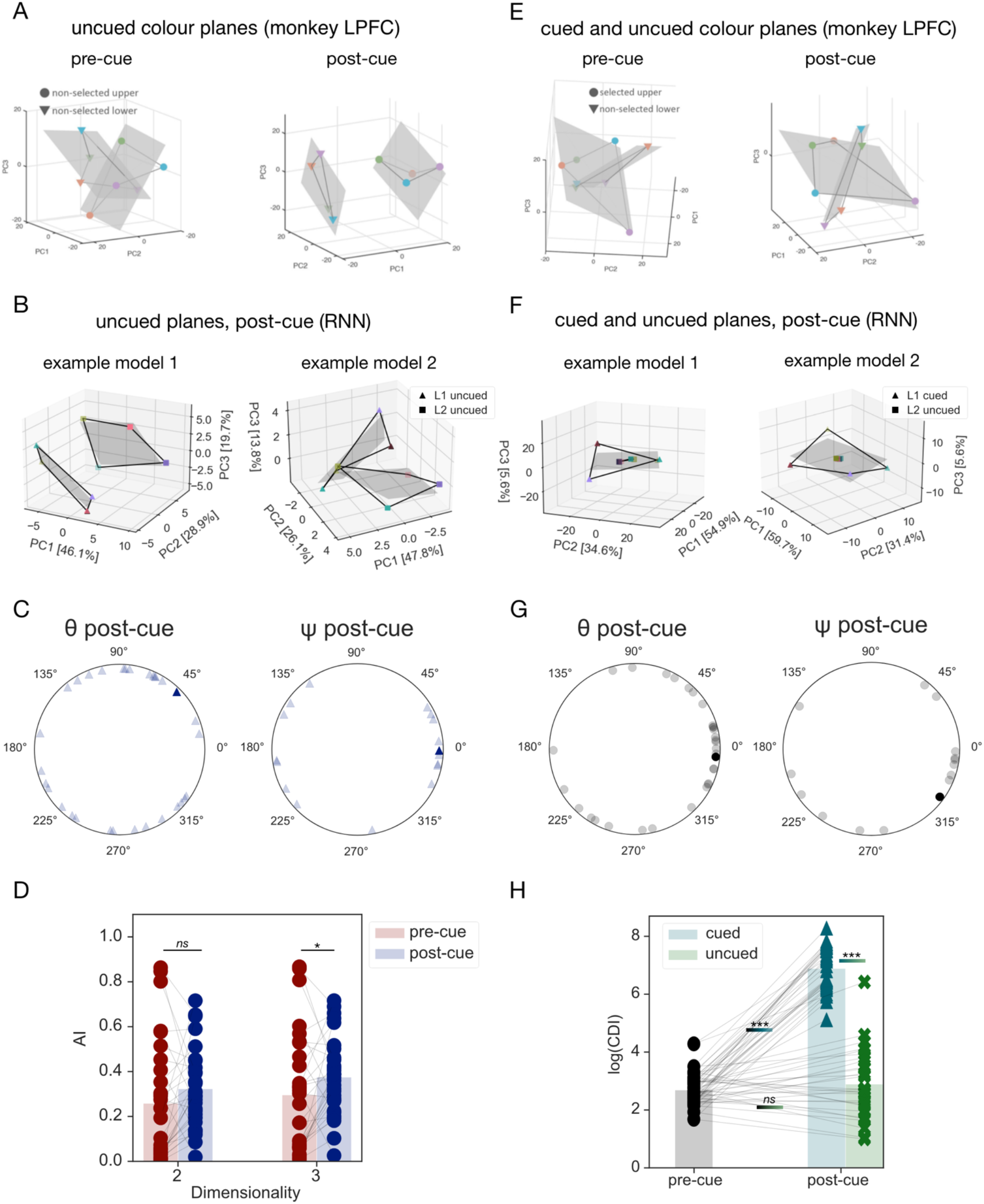
The geometry of the uncued item representations. **A**. Population activity patterns from macaque LPFC for the uncued items in the two delay intervals reported in [10]. Low-dimensional projections computed and visualised as described in Fig 2A. **B.** Hidden activity patterns for the uncued items, from two example models visualised in a 3-dimensional space. All conventions as in Fig 2B. **C.** Between-plane angles (*θ, left*) and phase alignment angles (*ψ, right*) between the two uncued colour planes in the post-cue delay. Individual models and population mean shown as transparent and opaque markers, respectively. **D.** Alignment index for the uncued subspaces in the pre- and post-cue delays. Note that prior to the presentation of the retro-cue, the coding format is location based. Therefore, the pre-cue data is the same as shown in Fig 2E. **E.** Projection of the population responses for cued upper and uncued lower items reported in [10]. **F.** Visualisation of the hidden activity patterns for cued and uncued items in the post-cue delay, on trials where L1 was cued. Data from two example models. Note the plane for L2 is severely compressed and thus hard to see. **G.** Angles between cued and uncued subspaces in the post-cue delay, averaged across the two retro-cue locations. *Left:* Between-plane angles *θ*, Right: phase-alignment angles ψ. Values for individual models shown in grey, average in black. **H.** Colour discriminability index (CDI) for the pre-cue subspaces (averaged across both locations, black circles) and cued and uncued subspaces in the post-cue delay (blue triangles and green crosses, respectively).

#### Representation of uncued items – *Cued/Uncued Geometry*

To successfully perform the task, the networks need to be able to prevent cross-interference between the two memory items (cued and uncued) across the entire trial duration, including the post-cue delay. In our next analysis, thus, we focussed on the relationship between cued and uncued items. We call this *Cued/Uncued Geometry* and note that this is different from analysing the relationship between each item (say, *colour-L1*) on those trials where it was cued relative to those where it is uncued. Instead, here we examine the relationship between *colour-L1* and *colour-L2* in the frame of reference of the cue.

It seems *a priori* plausible that, in order to minimise interference, the cued and uncued items are orthogonalized, as was observed in monkeys (**Fig 3E**). To test this prediction in our models, we calculated the angles between the cued and uncued subspaces and averaged them across the two trial types (defined by the location of the retrocue). Contrary to our expectation however, we found that the distribution of angles across the trained networks was not significantly clustered around 90 degrees (v-test: v(29) = −13.37, p < 1; **Fig 3G**, left). As this null finding could indicate that either (i) there was no significant clustering in the angles, or that (ii) the value around which they were clustered was different from our hypothesised mean, we followed it up by calculating the circular mean of the angles and performing a Rayleigh test. We found evidence for significant clustering in theta (z(29) = 6.02, p = .002) around the mean of 6.19°, indicating a parallel geometry. However, in contrast to the Cued Geometry in the post-cue delay, the Cued/Uncued Geometry did not permit a linear cross-readout of information between the cued and uncued planes over 50% of models formed mirror reflection or bowtie geometries, whilst for the remaining models, the cued and uncued planes were not significantly phase-aligned (v-test results: v(13) = 4.04, p = .064, N = 14; **Fig 3G**, right).

Furthermore, as was reported for monkey LPFC, the visualisation of the geometry in **Fig 3F** suggested that the uncued colour representations were much more compressed (i.e., less discriminable) in the representational space than their cued counterparts. To quantify this effect, we calculated the area of the polygons formed by the datapoints in the 3D space and compared this colour discriminability index (CDI) across conditions with contrasts based on the results reported for monkey LPFC [10]. We found that the mean colour discriminability in the post-cue delay was significantly higher for cued (CDI = 1294.1, arbitrary units) than uncued items (CDI = 45.46; one-sample t-test: *t*(29) = 48.95, *p* < .001, one-tailed; see also **Fig 3H**). Next, we tested the cued and uncued CDI against the pre-cue baseline. In accordance with previous reports from monkey LPFC [10] and human neuroimaging work [21,22], the cued CDI was significantly greater than the pre-cue baseline (CDI = 17.3; one-sample t-test: *t*(29) = 48.84, *p* < .001, one-tailed). For the uncued item, our hypothesis stated that there would be no significant increase from the baseline, as was reported in [10]. To test it, we performed a Bayesian non-parametric t-test. This returned a BF_10_ of 0.73, indicating anecdotal evidence for the alternative hypothesis of a difference in medians. For comparison, a two-tailed Wilcoxon signed-rank test was not significant (*W*(29) = 149, *p* = .085). These results suggest that to prevent interference between the cued and uncued items, the RNNs increase the representational discriminability of the former, whilst placing less importance on the geometrical arrangement of the Cued and Uncued planes at the end of the post-cue delay.

#### Learning dynamics and connectivity

Thus far, we have characterised the representational geometry formed by the RNNs and compared it to that observed in monkey LPFC. Next, we exploited the learning dynamics of the RNN to ask how the observed geometry emerged over the course of training.

Neural networks showed stereotypical learning dynamics characterised by a pronounced plateau period at approximately half of the initial loss value, buttressed by steeper declines both early and late (see **Fig 4A** for an example). We therefore decided to track the representational changes in the networks at three timepoints – at initialisation (*untrained*), the mid-learning plateau (*plateau*) and after convergence (*trained*). The timepoint used as the mid-training plateau was designated as the first local minimum in the derivative of the training loss with respect to time (for more details, refer to *Methods* and **S2 Fig**). We quantified the *Cued Geometry* across the different training stages by calculating the angle *θ* and the AI at the end points of the memory delays. The results of this analysis are shown in **Fig 4C**. *Untrained* networks propagated the orthogonal inputs throughout the pre-cue delay without changing their relative arrangement, as evidenced by an AI value of approximately 0 (**Fig 4C**, *top right*) and *θ* clustering around ±90°. This is expected, given that inputs to the network are orthogonal. After the presentation of the retrocue, both the AI and *θ* remained close to their pre-cue values. By the mid-training *plateau*, the networks had learnt to parallelise the two planes in the post-cue delay. Interestingly, however, this coding format also extended into the pre-cue delay, as evidenced by an AI value of close to 1 and *θ* clustering around 0°. As described in the previous section (*Representation of cued items*), the *trained* networks settled on an orthogonal plane configuration in the pre-cue delay, which was transformed into a parallel arrangement after the presentation of the retrocue. Thus, the network first learned to parallelise the inputs entirely, before gradually learning to keep them separate before the retrocue appeared.

**Fig 4.**
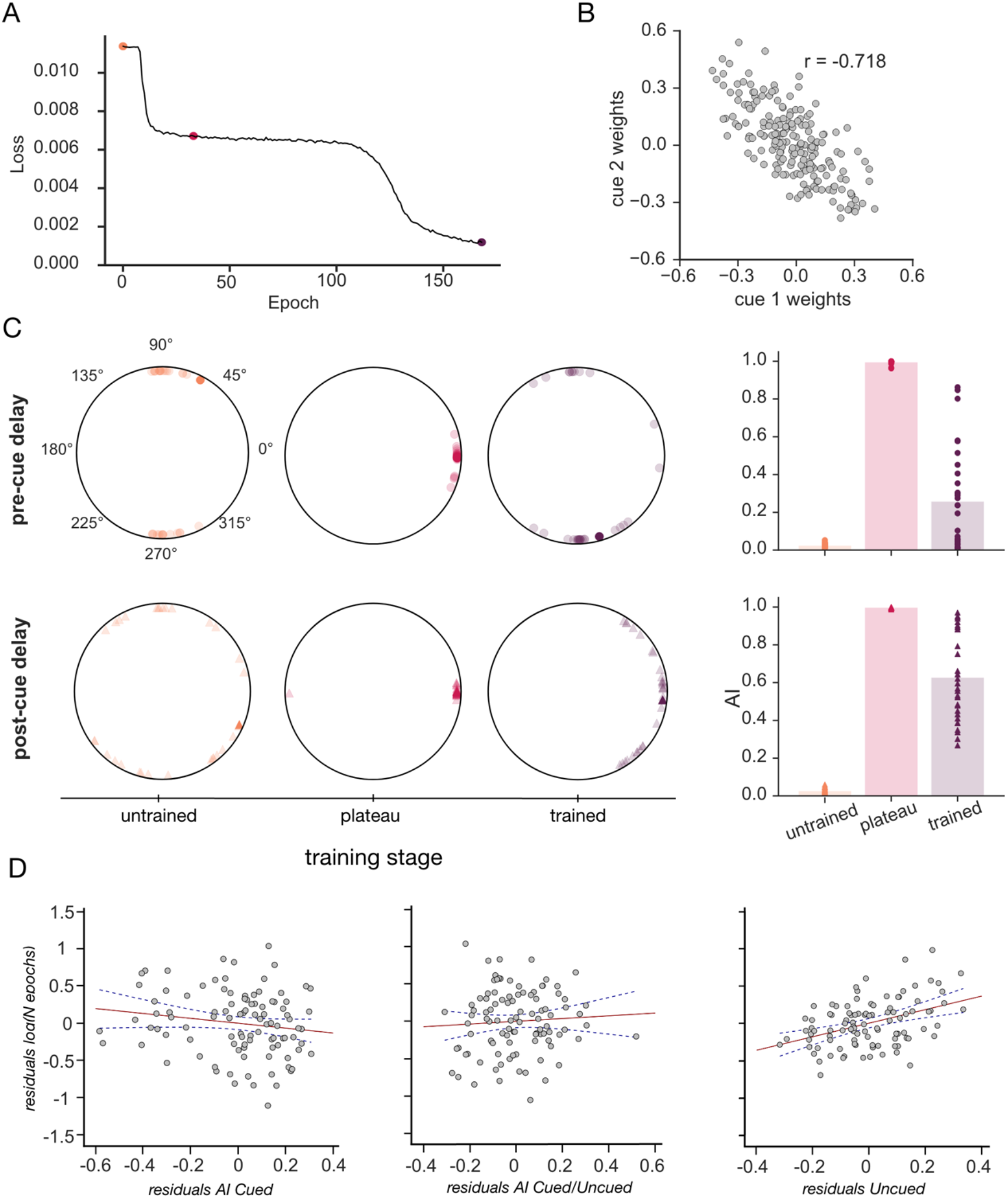
Learning dynamics. **A.** Training loss of an example RNN model plotted against training epochs. The three dots correspond to the timepoints chosen for the subsequent analysis: untrained (orange), plateau (magenta) and trained (purple) stages, respectively. **B.** Scatterplot of the weights between the two retro-cue input units and recurrent units for an example model. **C.** Comparison of the plane angles *θ* and AI between the cued planes across different training stages for the pre- (top row) and post-cue (bottom row) delays. *Left:* Plane angles at the untrained (orange), plateau (magenta) and trained (purple) stages. Values for individual models show in transparent colours, circular mean in opaque. *Right:* AI at the same training stages. Mean across models depicted as bars with individual datapoints overlaid. **D.** Partial plots showing the log of training epochs (y-axis) versus the three representational geometry metrics used as regression predictors (left to right: AI *Cued*,AI *Cued/Uncued* and AI *Uncued*). Values for individual networks shown as grey dots, regression lines of best fit in red alongside the 95% slope confidence intervals in navy dashed lines.

Next, we asked how the retro-cue triggered the representational changes observed in the recurrent layer. To address this, we analysed the connectivity patterns that were acquired over the course of training between the retrocue input units and the recurrent nodes. For each model, we calculated the correlation between the weights from each cue unit and the recurrent units (weights from an example model shown in **Fig 4B**). Consistent with other recent observations, we found that the models learnt opposing weight patterns for the two retro-cue inputs [11,23]. More specifically, the recurrent units that had positive connections weights with the first retro-cue unit had negative connection weights linking them with the other cue unit, and vice versa. Across all neural networks, this manifested as a significantly negative correlation coefficient (one sample t-test: *t*(29) = −68.17, *p* < .001). This negative correlation, in combination with the ReLU nonlinearities, ensures that hidden units that have inputs from one context units have none from the other and vice versa; this effectively creates two subpopulations of units which code for the context provided by the two retrocues.

Lastly, we investigated the relationship between the representational geometry observed in trained networks and their behavioural timecourse. To this end, we asked whether the orientation of the *Cued, Uncued* and *Cued/Uncued* subspaces (as quantified with the AI metric) was predictive of the training latency (time to convergence; see *Methods*) across all trained networks. To increase the power of this analysis (which unlike other analyses described here relies on correlations over the network population) we trained N = 100 models until the training loss fell below a common threshold value (see *Methods*). We then regressed the three AI values against the total number of training epochs. The regression model explained a significant proportion of the variance in training latency (*R^2^* = .28, *F*(3,96) = 12.29, *p* < .001). Out of the three predictor variables, to our surprise only the *Uncued Geometry* showed a significant and positive relationship with the number of training epochs (ß = 0.91, *t(96)* = 4.48, *p* < .001). This means that the more parallel the *Uncued geometry* in the post-cue period, the longer the network took to train. By contrast, the AI between the (i) *Cued* subspaces and (ii) *Cued/Uncued* subspaces did not contribute significantly to the model (*Cued:* ß = −0.125, *t(96)* = −0.374,*p* = .374; *Cued/Uncued:* ß = 0.17, *t(96)* = 0.878, *p* = .382). We were not expecting this result; which is further discussed below.

### Experiment 2: Exploring the effects of variable delay periods

In the simulations described thus far, we trained networks with a fixed number of delay cycles. In biological systems, maintenance processes can be successful even where the delay period length is unknown, and these conditions typically give rise to the formation of more stable dynamics (e.g. attractor states) in RNNs [24]. We thus asked whether (i) the networks could perform the task under novel delay period lengths and (ii) whether the representations they formed were dynamic or stable in nature, as well as how these aspects depended on whether the delay period during training was of variable or fixed duration. Lastly, (iii) we examined whether the geometry of cued item representations differed depending on the type of memory maintenance mechanism used by the network.

We sampled delay period durations randomly during training and test, so that networks could not learn to anticipate the precise delay length on any given trial. In this setting, we tested performance under three different conditions: a delay length previously experienced during training, as well as novel in- and out-of-training-range lengths. We found that networks trained with a fixed delay length could not perform the task with novel delay lengths – their performance, as measured by the mean absolute angular error rose from 16.37±0.94° (*M*±*SEM*) to 87.06±2.82° and 81.30±5.74° when evaluated on the 3 datasets, respectively (see also panel **A** in **S2 Fig**). In contrast, task performance in models trained with variable delay durations seemed to be less affected by the precise task timings (*M*±*SEM* = 7.72±0.22°, 36.46±1.94° and 18.60±2.01°, respectively). Note the overall performance difference between the fixed and variable delay networks is due a combination of different noise levels and training stop procedures used (for more details, refer to the *Methods* section).

Next, we studied the maintenance mechanisms learned by the networks trained with fixed and variable delay periods, using a cross-temporal generalisation approach. We trained LDA classifiers to discriminate between colour pairs using hidden activity patterns from a single timepoint and tested their performance on all the other trial timepoints (**Fig 5A** shows the cross-temporal generalisation scores averaged across models). We observed that neural networks trained with variable- and fixed-length delay periods both tended to maintain information across the memory delays using a temporally stable code. Accordingly, the mean off-diagonal delay cross-temporal generalisation scores were significantly higher than chance (0.5) in both groups (variable-length delay group: *M* = 0.62, one-sample t-test results: *t(29)* = 29.05, *p* < .001; fixed-length delay group: *M* = 0.59, one-sample t-test results: *t*(29) = 4.41, *p* < .001). However, as previously reported [24], the temporal stability of the delay-period code, as indexed by the ratio between the off- and on-diagonal delay decoding scores, was higher in the variable-than fixed-length delay group (*M* = 0.61 and 0.59, respectively) although this was only barely significant (independent sample t-test: t(58) = 1.71, *p* = .046, one-tailed).

**Fig 5.**
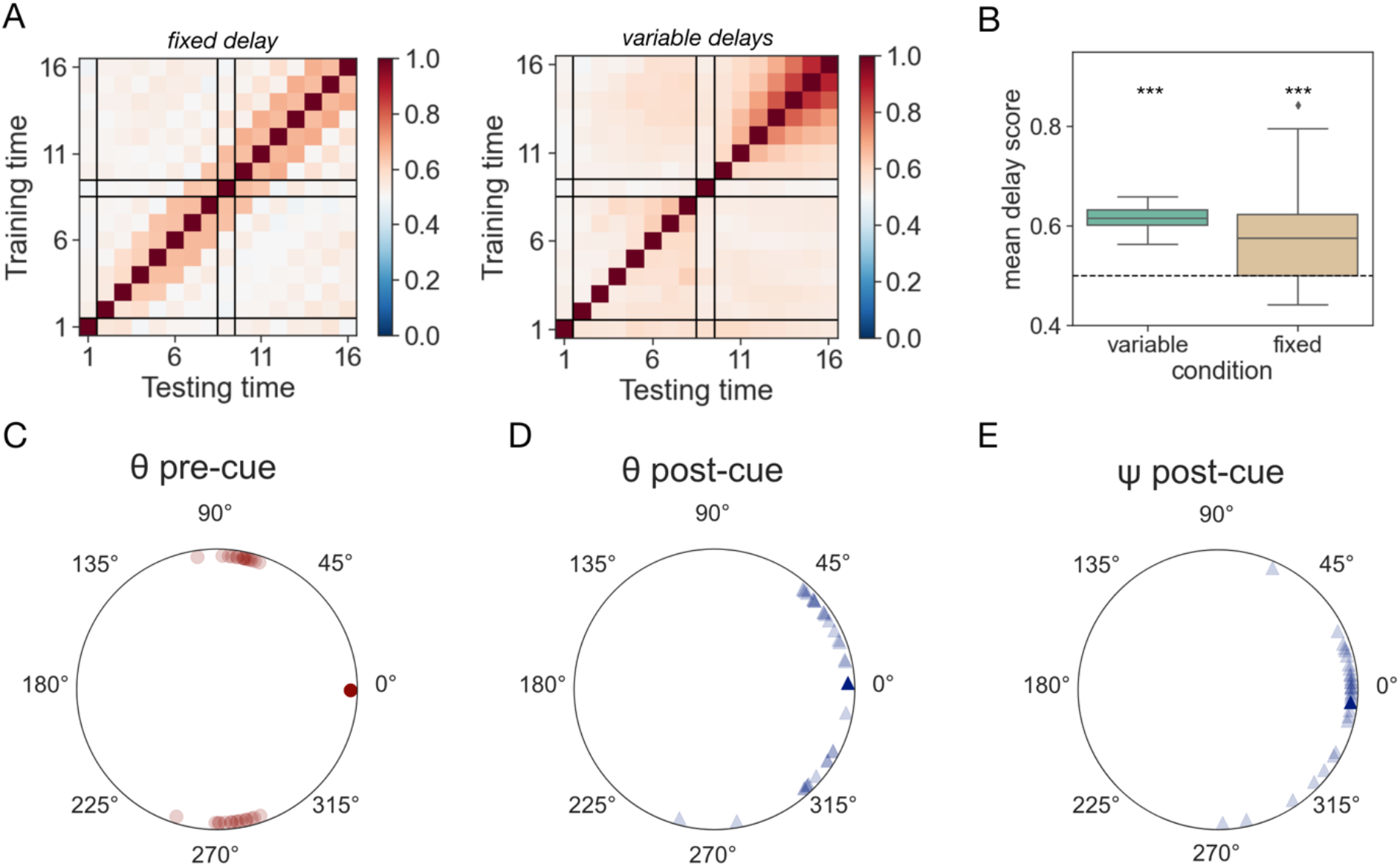
Maintenance mechanisms. **A.** Cross-temporal generalisation scores for decoders trained to discriminate between colour pairs. Data has been averaged across all trained networks. Black lines indicate the junctions between task events – stimulus presentation, pre-cue delay, retro-cue, and post-cue delay. *Left:* Average scores for models trained with a fixed-length delay interval. *Right*: analogous plot for models trained with variable delay lengths. **B.** Boxplots showing the distribution of mean cross-temporal decoding accuracy scores, averaged across the two memory delays, for all models. Variable delay interval length condition shown on the left in green, fixed delay length condition on the right in sand. Asterisks denote the results of the one-sided one-sample t-tests against the chance (50%) decoding level, *** corresponds to p < .001. **C.-D.** Plane angles *θ* between the cued subspaces in the pre- and post-cue delay periods, respectively. **E.** Phase alignment angles ψ between the cued subspaces in the post-cue delay.

Having confirmed that, compared to the fixed delay length networks, models trained with variable delay lengths learnt a task solution that generalised better across different timescales and showed more temporally stable memory maintenance dynamics, we analysed their hidden activity patterns to verify that our previous findings replicated under the new conditions. We focused on the *Cued Geometry* and observed that it matched the one previously described for the fixed-delay networks in Experiment 1. Across all trained models, the angles *θ* between the two cued subspaces were significantly clustered around the 90° mark in the pre-cue delay (v-test results: *v*(29) = 27.7, *p* < .001) and 0° in the post-cue delay (circular mean = 2.73°; *v*(29) = 22.27, *p* < .001, see also **Fig 5C**). The mean absolute difference between pre- and post-cue angles was also significantly different from 0° (one-sample test for mean angle, *M* = 103.34°, 95%*CI* = 75.39°, 131.28°). Furthermore, the Cued planes were also significantly phase-aligned in the post-cue delay (mean ψ = −5.44°; v-test results: *v*(28) = 25.09, *p* < .001; note one model was excluded from this analysis due to forming a mirror image geometry with *θ* = −105.21°) . This pattern of results was further confirmed by the AI analysis. The mean pre-cue AI (AI = 0.09 and 0.11 for 2 and 3 dimensions, respectively) was significantly lower than the mean post-cue AI (AI = 0.13 and 0.14, respectively) in both target dimensionalities (one-tailed one-sample t-test: *t*(29) = 4.72, *p* < .001; *t*(29) = 2.94, *p* = .003). We note that although the differences between pre- and post-cue delays are qualitatively similar, these values are overall lower (i.e., planes are more orthogonal) than for Experiment 1.

### Experiment 3: Modelling the effect of retro-cue validity

Behavioural performance on multi-item working memory tasks is enhanced when participants are presented with a valid retro-cue, correctly informing them about the item they will be probed on, relative to retro-cues that are invalid (i.e., point to one of the unprobed memory items) or neutral [7,8]. The precise mechanism by which this effect arises is unclear. One account, consistent with the results described above (and those in monkeys) proposes that the presentation of the probabilistic retro-cue triggers a transformation of the prioritised item into a new, response-oriented format, thus increasing the robustness of its representation [9].

Seeking a normative account of this phenomenon, we trained RNN models on a modified version of the task from Experiment 1 (see **Fig 5E** for a schematic of an example trial). As before, the networks were first presented with the two coloured stimuli, followed by a delay, retro-cue, and another delay. However, unlike in the experiments conducted thus far, the cue indicated which item was trial-relevant with varying degree of validity and was followed by a deterministic probe and another memory delay. We manipulated the percentage of training trials on which the retro-cue and probe inputs matched (*retro-cue validity*). We designed 2 experimental conditions of interest, where the retro-cue validity was set to 75 and 50%, in addition to a control condition with validity equal to 100% (this latter experiment resembles those in Exp.1, except for differences in timing of the trial events). As for other experiments (except where indicated), we trained 30 models per group.

When retro-cue validity experienced during training was 75%, model performance (as measured by MSE recall) was higher for validly than invalidly cued trials (see **Fig 6B**). In contrast, when the information provided by the retro-cue was maximally ambiguous (validity = 50%), performance on valid and invalid trials was matched (note that in this case, even though the retro-cue signal does not correlate with the trial-relevant item, it is still possible to make a distinction between valid and invalid trials, as the identities of both the retro-cue and probe were signalled using the same pair of input nodes, valid trials refer to those on which the same node was activated twice). To understand these effects, we leveraged the fact that errors on delayed-estimation tasks such as the retro-cueing task are thought to arise from a combination of imprecise reports of the target item, incorrect reports of the non-target item, and random guesses. Adopting an approach familiar from human behavioural studies, we fit a mixture model incorporating these error terms to the individual model choices [25]. The model had four free parameters, corresponding to memory precision (*K*), probability of recalling the target item (*pT*), probability of recalling the non-target item (*pNT*), and probability of making random guesses (pU). We compared each fitted parameter type across conditions (validity = 75% and 50%) and trial types (*valid, invalid*) with a 2×2 mixed model ANOVA (or a non-parametric equivalent in the case of the probability parameters, which had highly unequal variances between conditions). To summarise the results below, we found that four all four parameters tested, there was both a main effect of trial type and an interaction between trial type and condition, suggesting that the superior performance of the neural networks on validly cued trials under probabilistic conditions can be accounted for by precision, recall, protection against intrusions of the uncued item, and nonspecific lapses. To avoid cluttering the text, we report full statistics for these comparisons in a supplementary note (**S4 Note**), but they are also visible in **Fig. 6C** below.

**Fig 6.**
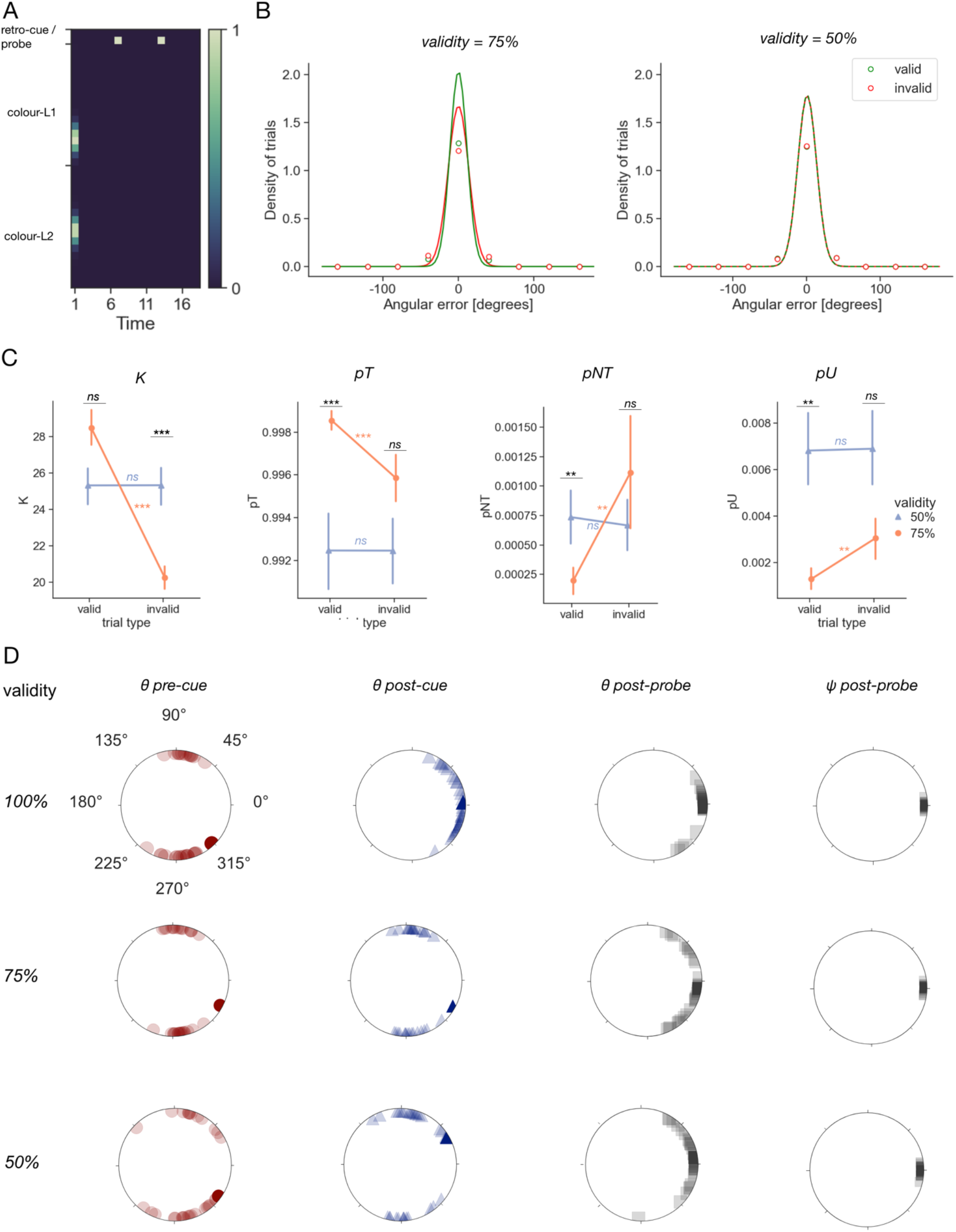
Probabilistic cues. **A.** Task structure. **B.** Distribution of errors made by the models after convergence (dots and spikes, *M*±*SEM*) shown with a best-fit von Mises pdf (line). Performance on trials where retro-cue matched the probe (*valid*) shown in green, mismatch (*invalid*) trials shown in red. Data plotted for models trained under 2 different cue validity conditions −75 and 50%. **C.** Comparison of the mixture model parameters (left to right panels: *K, pT, pNT* and *pU*) across cue validity conditions (50 and 75%, corresponding to lavender triangles and orange circles, respectively) and trial types (valid, invalid; shown on the x-axis). Asterisks denote the significance levels of the post-hoc tests described in the main text: ***: *p* < .001, **: *p* < .01, *ns:p*>= .05. Comparisons within a validity level shown in orange and lavender, comparisons across different validity levels (but for the same trial type) denoted in black. **D.** Distribution of plane angles *θ* and phase-alignment angles ψ formed in the delay intervals for all trained networks. Values for the pre-cue, post-cue and post-probe delays shown as red circles, blue triangles and grey squares. Data from individual models shown in opaque, and grand averages in solid colours. Plots for models trained under trained under the deterministic (retrocue validity = 100%) and non-deterministic retrocue validity conditions (75% and 50%) ordered top to bottom rows, respectively.

Having explored the nature of the behavioural effect observed in **Fig 6B**, we then compared the neural geometries formed in networks trained under different retro-cue validity conditions. Once again, we were interested in the relationship between the trial-relevant item subspaces in the different delay periods. We found that networks trained under the deterministic regime (where the retro-cue was valid on 100% of trials) kept the information in orthogonal subspaces during the pre-cue delay (v-test: *v*(29) = 25.79, *p* < .001) and underwent a configurational change into parallel planes after the retro-cue presentation (v-test: *v*(29) = 25.09, *p* < .001) that prevailed into the post-probe delay interval (v-test: *v*(29) = 28.16, *p* < .001; see also **Fig 6D**). The post-probe planes were also significantly phase-aligned (v-test: *v*(29) = 29.96, *p* < .001). In contrast, in the networks trained under non-deterministic conditions (cue validity 75% and 50%), the retro-cue did not trigger this representational change (v-test: *v*(29) = 3.45, *p* =.186 and v(29) = 5.34, *p* = .084; respectively). Under these conditions, the information was transformed into the parallel format later in the trial, in the post-probe delay (*v*(29) = 25.49, *p* < .001 and *v*(29) = 25.49,*p* < .001; respectively). At that timepoint, the cued planes were also phase-aligned (*v*(29) = 29.96, *p* < .001 and *v*(28) = 28.92, *p* < .001; for cue validity 75% and 50%, respectively).

Finally, we asked whether the representational geometry in the post-probe period developed by the trained models was behaviourally relevant. Like previously described for Experiment 1, we regressed the AI metrics for the *Cued, Uncued* and *Cued/Uncued* geometries against the total number of training epochs completed by the networks, separately for each validity condition. The results of this analysis largely followed the same pattern as described for Experiment 1. In the 100% validity condition, the regression model explained a significant proportion of the variance in training speed across networks (*R^2^* = .23, *F*(3,96) = 9.49, *p* < .001). The AI between the *Uncued* subspaces was positively predictive of the number of training epochs (ß = 4.59 × 10^-4^, *t(96)* = 4.71, *p* < .001), providing a separate replication of this effect. Additionally, the AI between the *Cued* subspaces bore a significant negative relationship with the training speed (ß = −8.93 × 10^-4^, *t(96)* = −3.05, *p* = .003), whilst the AI between the *Cued/Uncued* subspaces did not contribute significantly to the model (ß = 5.51 × 10^-5^, *t(96)* = 0.52, *p* = .602). The same pattern of results was observed in the 75%(*R^2^* = .42, *F*(3,96) = 23.03,*p* < .001; AI *Uncued:*ß = 0.006*, t(96*)= 4.88,*p* < .001; AI *Cued:*ß = −0.004*, t(96*)= −3.65,*p* < .001; AI *Cued/Uncued:*ß = 0.004, *t(96)* = 2.07, *p* = .041) and 50% (*R^2^* = .38, *F*(3,96) = 19.46, *p* < .001; AI *Uncued:*ß =0.046*, t(96*)= 5.52, *p* < .001; AI *Cued:* ß = −0.024, *t(96)* = −2.93, *p* = .004; AI *Cued/Uncued:* ß = 7.88 × 10^-4^, *t(96)* = 0.07, *p* = .947) validity conditions (see also **S3 Fig**). In other words, we see reasonable convergence between results obtained in these probabilistic conditions and the deterministic conditions explored by [10].

### Experiment 4: Parallel geometry for more robust maintenance

Thus far, we have observed that RNNs trained on variants of the retro-cueing task learned to transform the cued item information into a common frame of reference after the retrocue (or probe, in the case of probabilistic cues). Whilst this representational format affords the re-use of a common readout mechanism across all trials (i.e., irrespective of the retro-cue identity), it is not the only solution to the task. In fact, we observed that in the probabilistic paradigm described above, networks trained with no post-probe delay did not develop this geometry, and instead maintained the cued item information in orthogonal planes throughout the entire trial (data not shown). Given this observation, we reasoned that the parallel plane format might be beneficial when there is a pressure to maintain the cued information in memory after the retro-cue [9]. To test this hypothesis, we trained models (as before, N = 30 per task variant) on variants of the original task from Experiment 1 (with a deterministic retro-cue and no probe), except that now we parametrically varied the length of the post-cue delay interval, whilst holding the overall trial length constant (see **Fig 7A**). We then calculated the AI between the cued subspaces prior to and post retro-cue presentation and compared them between the different task variants (**Fig 7B**, red dots and line). We found that, in agreement with our previous results, all models maintained the cued item information in orthogonal subspaces before the retro-cue presentation, regardless of the task timings.

**Fig 7.**
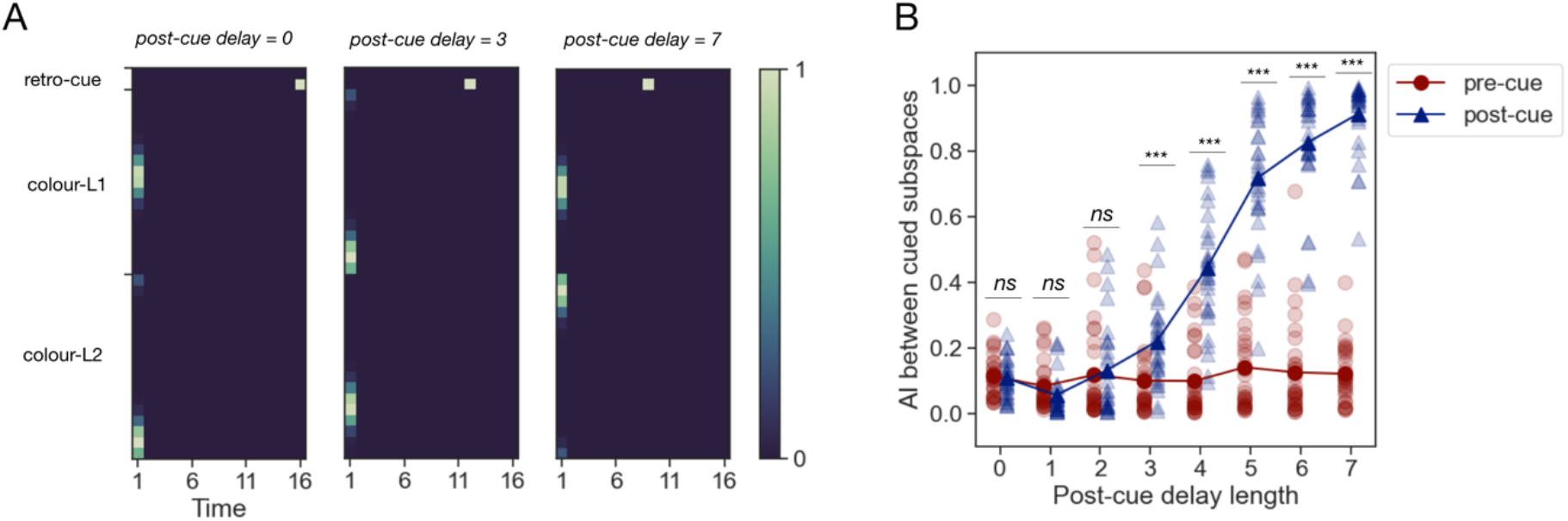
Comparison of cued item geometry across networks trained under various post-cue maintenance pressure conditions. **A.** Example trials for a subset of the parametric family of retro-cueing tasks used. Overall trial length was constant, and the ratio between pre- and post-cue delay lengths was varied. Conventions as in Fig 1B. **B.** Comparison of cued subspace AI during the pre- (red circles) and post-cue (blue triangles) intervals between networks trained with different post-cue delay lengths. Asterisks denote the significance levels of the contrasts described in the main text: ***: *p* < .001, **: *p* < .01, *ns*: *p* >= .05.

However, only models trained with longer post-cue delay lengths underwent the previously noted transition into a parallel subspace format prior to producing a response, whilst networks trained with shorter delays learnt to read-out cued colour information from the low-dimensional orthogonal subspaces (**Fig 7B**, blue dots and line). To test the statistical significance of this finding, we compared the pre-to post-cue difference in AI with a Kruskal-Wallis test, which showed a significant difference in this metric across the different post-cue delay length conditions (*H*(7) = 199.7, *p* < .001). We then followed up with custom contrasts comparing the pre- and post-cue AI at each level of the post-cue delay length factor. This revealed no significant differences with delay lengths below 3 cycles (*t*(232) = 0.27, *p* = .789; *t*(232) = −1.16, *p* = .249; *t*(232) = −.61, *p* = .546, in increasing order) and a significantly higher post-cue AI otherwise (*t*(232) = 5.04, *p* < .001; *t*(232) = 14.53, *p* < .001; *t*(232) = 24.3, *p* < .001, *t*(232)= 29.52, *p* < .001, *t*(232) = 33.36,*p* < .001, in increasing order). Note that the degrees of freedom here are computed from the ANOVA (8 conditions × 30 models = 240 datapoints, see methods).

## Discussion

In this study, we describe a detailed study of the short-term memory representations that arise in RNNs during a retro-cueing task. We report that these are similar to those previously described in [10] for monkeys. In macaque LPFC, it was reported that prior to the presentation of a retrocue, information about individual memory items was maintained in short term memory along orthogonal axes of the neural state space. In contrast, following the retrocue, representations of the prioritised items were rotated into a common subspace. As seen across the four experiments reported in this paper, the RNN reliably captured this orthogonal-to-parallel transformation when the relationship between the retrocue and the trial-relevant item was deterministic (but not when it was probabilistic, see Exp.3). Building on this representational correspondence, we then characterised how and why the described geometry emerged, in addition to revealing its mechanistic basis, thus providing testable predictions for future neurobiological studies.

At first glance, the orthogonal geometry in the pre-cue delay might seem a trivial finding, given that the information about the two stimuli is supplied to the RNN via independent (and uncorrelated) input channels. Accordingly, this geometry is present even in naïve networks (**Fig 4C**). However, during the pre-cue delay, over the course of training the networks first learn to arrange both mnemonic representations into parallel neural subspaces before converging back onto the orthogonal geometry. This coding format seems well suited to satisfy the computational demand of the task in the pre-cue delay – namely, the minimalization of interference between multiple memory items. This interpretation also fits well with other findings from the literature. For example, a recent sequential working memory study reported that information about individual stimuli from a sequence was also maintained in orthogonal neural subspaces in macaque LPFC [15]. Similar observations have also been made in other brain regions, species and task [12,13,27] as well as in neural networks [23,26,27], suggesting that factorising representations into orthogonal subspaces to prevent cross-interference is a general coding principle shared by both biological and artificial networks.

Interestingly, the principal mechanism for preventing interference between the cued and uncued items in the post-cue delay in the RNNs seems to be the modulation of the discriminability of the cued and uncued stimuli. In the representational geometry framework, this corresponds to the overall size of the colour rings. In agreement with previous reports from both macaque [10] and human research [21,22], we found evidence for an increase in colour discriminabillity for the cued item. With this mechanism in place, the geometrical arrangement of the two planes should not bear much relevance and might be difficult to estimate – accordingly, we found the angles between the cued and uncued planes to be roughly centered on 0 degrees, albeit with a considerable variability across models (in comparison to the *Cued Geometry* angles).

In the post-cue delay interval, the representations of cued memory items were arranged into parallel subspaces in both the monkey LPFC [10] and the RNN. This geometry abstracts over the now-irrelevant stimulus location and aligns the colour representations into a common template space. Consequently, the same downstream readout mechanism can be used to generate a response, regardless of the retrocue identity. Interestingly, the authors in [10] also report the same neural geometry in a related task where the cue is provided prior to the stimulus presentation, at the onset of the trial. Whilst we did not model this task variant, we predict that the neural geometry developed by the RNNs under such conditions would similarly match. This prediction is based on observations of parallel low-dimensional manifolds in other tasks incorporating a response generalisation element, which have been made in both biological and artificial networks [16–19]. Such findings collectively imply that the parallel geometry might too be a general coding principle, its function being to facilitate the generalisation of knowledge across different behavioural contexts.

Nor does it seem like the parallel plane geometry in the post-cue delay is a trivial consequence of the way we set up the task. In Experiment 4 we show that this geometry only emerges when the cued information must be maintained in memory for several cycles prior to the response. In contrast, when there is only a short post-cue delay, the RNNs continue to maintain the cued information in the orthogonal format adopted in the pre-cue delay period. Thus, our results suggest that the parallel geometry might additionally confer a robustness to working memory maintenance, providing a normative explanation for its emergence in the context of the retro-cueing task, as well as a testable prediction for future neurobiological studies. We note that these speculations, based on our simulations, could be confirmed with exact analysis of neural network dynamics (where these become available).

Whilst the *Cued* and *Cued/Uncued* geometries are interpretable with respect to the task demands as outlined above, this is not the case for the *Uncued* geometry. We found the *Uncued* geometry to be variable between the different network replicants. Surprisingly however, this geometry was also highly predictive of network behaviour – in both Experiments 1 and 3, it proved to be a reliable predictor of the training latency. This relationship was negative, indicating that to reach the same level of performance, models that relied on a more parallel *Uncued* geometry needed to complete more training iterations than those that orthogonalised the uncued item subspaces. We interpret this result to be a consequence of using random network initialisation, which would predispose some networks to a more parallel *Uncued* geometry even before training commenced. If the uncued planes happened to be aligned with the readout subspace in a network, it would be more prone to responding based on the uncued item and thus need more training iterations to overcome this response bias. It is unclear whether this geometry bore any relationship to behaviour of the monkeys reported in [10]. Interestingly however, the authors reported a negative correlation between the degree of the *Cued* plane alignment and the magnitude of behavioural error. In agreement with their report, we found it to be positively predictive of the training latency in Experiment 3, signifying a performance advantage in networks characterised by a more parallel geometry. However, this effect was not detectable in Experiment 1. We are thus not entirely convinced by this explanation.

The similar representational geometry with the macaque suggests that the LPFC might use a similar computational principle to solve the retro-cueing task. Thus, in later experiments we capitalised on the inherent accessibility of the RNN model and extended our analyses to two aspects not explored in [10] – the (i) emergence and (ii) mechanistic basis of the reported representational geometry. As artificial neural network models are not endowed with representational schemes by hand and instead develop them through training, they offer a normative account of how representational properties can be acquired through learning in biological brains. To this end, we provide a description of the stereotypical representational geometries the RNNs develop at different stages of training. The existence of such learning transitions can be verified in vivo in future studies, to ascertain whether biological learning follows similar principles to the learning algorithms used for training RNNs.

Our final contribution lies in the exploration of a related paradigm often used in behavioural experiments, whereby the relationship between the retrocue and the trial-relevant item is probabilistic. In Experiment 3, we recreated the key behavioural results characteristic of human participants performing the task. We found that, under conditions where the cue was on average valid (75% validity paradigm), recall performance was improved on trials where the retrocue correctly pointed to the to-be-probed item and degraded otherwise, relative to the corresponding performance levels in a fully probabilistic set-up (50% validity paradigm). These effects have also been reported in human participants performing equivalent tasks (for a review see [28]). In a further parallel to the human literature, we found the locus of the retrocue benefit to lie within an increased probability of recalling the target item [29–36] and reduced probability of making swap or random errors [33]. However, we did not find a reliable increase in the target item recall precision on valid trials, which has been described in some human studies [30,33–36]. This discrepancy most likely reflects the training procedures followed, which lead to matching overall performance levels on valid trials in both the 75% and 50% validity regimes.

To our surprise, only networks trained with 100% valid retrocues were characterised by a parallel *Cued* geometry in the post-cue delay interval. In contrast, networks trained under 75% and 50 validity conditions continued to maintain the cued item information in orthogonal subspaces until the post-probe delay, when the planes were rotated into a shared subspace. This finding was particularly surprising in the case of the 75% validity condition, given that the presence of the behavioural effect described above strongly suggested that the networks relied on the retrocue input to solve the task. Thus, it appears that a parallel post-cue *Cued* geometry forms part of an optimal task solution only when the retrocue is fully informative about the trial-relevant item, however the reasons for it are unclear.

Overall, our findings are consistent with theories proposing that retrocues benefit performance partly by triggering a representational transformation into an action-optimised format [37]. We also find evidence for the attentional strengthening hypothesis of the retro0cueing effect [28] – in Experiment 1, we observed a selective increase in colour information for the cued item in the post-cue delay, in accordance with other observations from primate recordings [10] and human neuroimaging [21,22]. On the other hand, our findings are inconsistent with accounts proposing the removal of uncued items post-cue [28] – in Experiment 1, we found that the post-cue delay was still characterised by significant information about the uncued item. The same result was reported in monkey LPFC [10]. On a broader scale, this work speaks to the utility of the representational geometry approach for revealing computational mechanism in both artificial and biological networks. As seen with regards to the *Cued* geometry, characterising the geometric structure of the stimulus representations and its transformations across different stages of processing permits inferences about the nature of the computations learnt by the network. Over and above that, this framework affords direct cross-modal comparisons within a context of a given paradigm, as well as can shed light on computational similarities across different task domains.

## Methods

### Model architecture

Each recurrent neural network (RNN) consisted of an input, recurrent and output layer. The input layer consisted of 36 units in total. There were 2 binary units corresponding to the location retro-cues (and probes, if used), 17 colour units encoding the feature value of the stimulus shown at location 1 (L1) and another set of 17 units encoding the feature value at location 2 (L2). Here, we assume that feature values are hues drawn from a circular colour space, but our simulations hold for any circular feature space (e.g., orientation).

Each of the feature-sensitive units had a circular normal tuning function centred on one of 17 equally spaced values. On stimulation, each unit *i* produced activity *u_i_* which depended on its preferred colour ϕ and the current stimulus colour *α*, and was given by the von Mises probability density function (1.1):

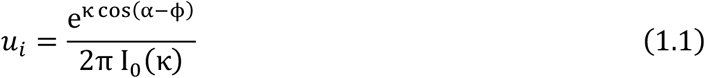

where I_o_(κ) is the modified Bessel function of order 0, and κ corresponds to the tuning width (chosen to be 5.0 for the reported simulations). The distribution of input activity was rescaled to be in the range [0,1]. The retrocue inputs (which occurred later) were one-hot signals denoting whether the top or bottom cue was presented.

The recurrent layer of the network consisted of *N_rec_* = 200 units. The activity of the hidden unit vector *h_t_* at time-step *t* was given by:

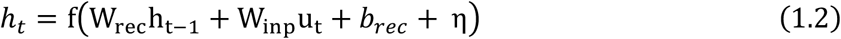

where W_rec_ and W_inp_ correspond to the recurrent and input weight matrices, u_t_ is the vector of input unit activity on timestep t, *b_rec_* is a bias term and η is a noise term drawn from N(0, σ^2^). At the beginning of each trial, the hidden state was initialised at 0. The activation function f used was the rectified linear unit (ReLU) function. On the final timestep, hidden activity was read out by a single layer of N_out_ = 17 units, each associated with a response probability 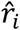:

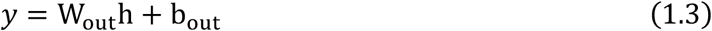

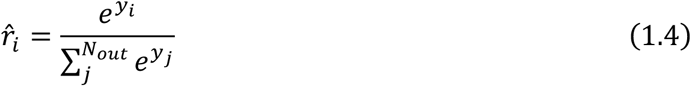

where W_out_ is the output weight matrix and b_out_is a bias term. The resulting output *y* is transformed to vector of responses probabilities (of making each of the N_out_ = 17 colour choices) by a softmax function (1.4).

### Experiment 1

#### Task structure and training procedures

Each trial started with the presentation of two colour-location stimuli for one cycle, followed by 7 cycles of (*pre-cue*) delay, a single-cycle retro-cue input, and another 7-cycle (*post-cue*) delay. Networks (N=30) were trained on a dataset consisting of 512 unique trials, corresponding to all the possible combinations of 16 colours at the two stimulus locations and 2 retro-cues. We initialised the recurrent weights as an orthogonal matrix [38] and used the standard Xavier initialisation for all other learnable network weights and biases. Each trial was associated with scalar feature values (*α_cued_* and *α_uncued_*), corresponding to the value of the tuning curve peak in colour space for the cued and uncued stimuli. Network parameters were tuned with stochastic gradient descent (batch size = 1 trial) using the RMSprop optimiser. We defined the ground truth response *r* as a one-hot vector over output units, with a 1 at α_*cued*_ and zeros elsewhere. The loss term was given by the mean squared product of the (i) distance between *r* and the network output 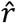 and (ii) the circular distance between the output unit tuning centres ϕ and α_*cued*_:

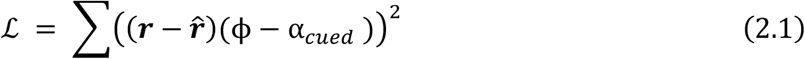

Note the latter term ensures that the penalty the network receives is proportional to the circular distance from the correct response, and thus constrains the output values in 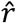 to have a smooth distribution with a peak at the correct answer location.

We conducted a hyperparameter search over a range of different recurrent layer sizes, learning rate values and noise levels. The data reported in this paper comes from networks with 200 recurrent units and trained with learning rate set to 0.0001. We chose σ = 0.07, as it resulted in model performance that was qualitatively similar to that reported for monkeys (measured by the absolute mean of the error distribution). The pattern of results did not differ between networks trained under different hyperparameter choices. We stopped model training as soon as the loss function reached its final plateau. This moment was defined by two criteria: the (negative) slope of the loss function over the last 15 epochs falling below a threshold value of 2e-05, and the mean loss value over the most recent epoch falling below a threshold of 0.0036 (a value that corresponded to the performance level, as measured by absolute mean angular error, reported in monkeys). Note that imposing a threshold on the absolute loss value was necessary to avoid stopping training prematurely (i.e., at the mid-training learning plateau, see also **S1 Fig**).

#### Data analysis

We evaluated the trained models on the complete training dataset after freezing the weights (i.e., disallowing further gradient updates). The testing dataset contained 100 repetitions for each of the 512 unique trials (note that this was not redundant because the noise term introduced variability). To generate network responses, we sampled colour choices proportionally to their probabilities as given by the output layer activations (see equation 1.3).

#### Plane fitting, angles and phase alignment

To obtain planes of best fit for each stimulus location, we adapted the method described in [10]. In short, for each trained RNN, we sorted the trials into B = 4 colour category bins and L = 2 locations, giving a total of B × L = 8 conditions. For each timestep of interest (the end points of the memory delays), we then calculated the mean hidden unit activity *h* by averaging within these bins, giving rise to a population activity matrix **X**_*full*_ **of size** 8 × *N_rec_*. We then applied principal component analysis (PCA) to the data in two steps: first, reducing the dimensionality of X^*full*^ to the reduced-rank matrix *X^3D^* **of** size 8 × 3; and second, reducing each location L separately (corresponding to a 4 × 3 input subset) to a 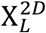 matrix of size 4 × 2. The two principal components (PCs) calculated during the second step correspond to the directions of maximum variance in the data, and thus define the axes of the best-fit planes for the given location in the input (3D) space.

To calculate the angle θ between the two planes, we first applied a correction to the plane-defining vectors for both locations that put them in a common frame of reference. This was necessary because the vectors obtained from PCA correspond to the directions of maximum spread in the data rather than, for example, specific sides of the quadrilateral defined by the datapoints. If the directions of maximum variance differ between the two locations, this could influence the value of the angle between planes calculated. For example, imagine the colour-category datapoints form rectangles for both stimulus locations considered. However, the longer side for the rectangle in L1 corresponds to the line joining the ‘yellow’ and ‘red’ datapoints, whereas for L2, this is the shorter side of the rectangle. In this case, despite the data rectangles being parallel (as well as phase-aligned), the sets of plane-defining vectors obtained from PCA for both locations would differ in the ordering of the calculated vectors. As the angle between planes is based on calculating the plane using the cross-product (see equation 5.1), which is sensitive to the vector order, this would then result in the angle taking on a value of 180°rather than 0°. Thus, to make the metric insensitive to any distortions to the square geometry, we applied a correction to the set of vectors obtained from PCA. The first vector from the set was rotated to be collinear to a side of the data quadrilateral (giving rise to PC1_L_corrected__), and the second vector was rotated by the shortest angular distance to be orthogonal to the former (PC2_L_corrected__). After applying this correction, we calculated the normal n_L_ for each location L:

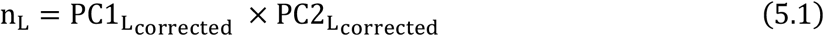

The angle between the two planes was then be defined as:

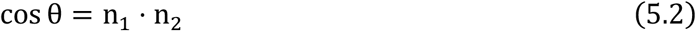

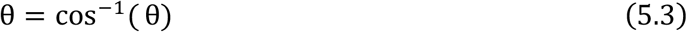

Note that following the correction, an angle of 0°is only possible when the two planes are parallel and are rigid rotations and/or translations of one another. In contrast, parallel planes that are mirror reflections of one another and bowtie geometries would result in an angle of 180°.

To test for significant clustering of the plane angles, we used Rayleigh, v- and circular one-sample mean tests. All tests assume a null uniform distribution around the entire circle, but the range of cos^-1^ is restricted to [0,180°]. Therefore, we set the sign of each angle as the sign of the z-coordinate of ni in the 3-dimensional coordinate system obtained from the PCA analysis, thus extending the range to the full [−180°,180°] interval. Furthermore, all tests also assume a unimodal distribution as the alternate hypothesis. Therefore, in cases where the data distribution was bimodal (i.e. pre-cue angles, as well as the pre-post angle difference), we first rectified the plane angles (to force the distribution to be unimodal) and then multiplied the individual values by a factor of 2 (to extend the possible range to [0,360°]).

Phase alignment was defined as the rotation angle ψ that minimised the distances between corresponding points from the two locations. Planes with absolute θ values greater than 90° (which correspond to mirror reflections) were excluded from the analysis, as were bowtie geometries. For all other geometries, we projected the entries from X^*3D*^ onto their corresponding planes, before forcing the planes to be co-planar by rotating plane 2 by -θ°. We then applied the orthogonal Procrustes analysis. This method finds an orthogonal matrix R which most closely maps one matrix onto another (in our case, the sets of points corresponding to locations 1 and 2). Phase alignment angle ψ was defined as:

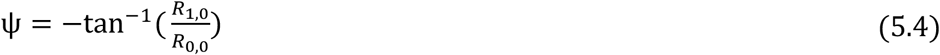

To assess for significant phase alignment between planes, we performed a v-test with the hypothesised mean set to 0°.

#### Subspace alignment index (AI)

The alignment index between a pair of subspaces was calculated as described in [20]. In short, we defined two matrices, S_1_ and S_2_, containing the binned activation patterns corresponding to the subspaces of interest. For example, when calculating the AI between the L1 and L2 subspaces during the pre-cue delay, the two matrices would correspond to the first and last B rows of **X***^full^*. Next, we calculated the covariance matrices for S_1_ and S_2_ and performed eigenvalue decomposition. The alignment index between S_1_ and S_2_ is then given by:

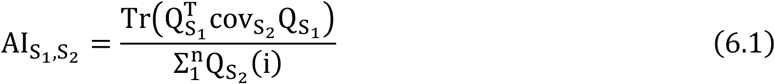

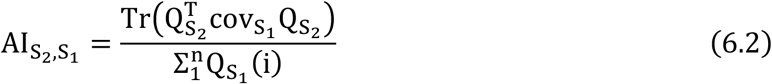

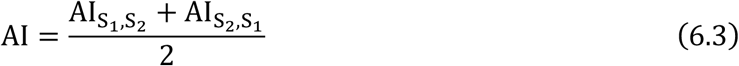

where cov_S_1__ corresponds to the covariance matrix of S_1_, and Q_S_1__ is the matrix containing its eigenvectors.

For the cross-validated AI analysis of *Cued Geometry*, we used half of the data to calculate the AI values for the L1 subspaces between the two memory delays, repeated this step for the L2 subspaces, and compared the two values to define planes as *unrotated* and *rotated* according to their AI values. Using the other half of the dataset, we then once again calculated the AI, this time comparing the *unrotated* and *rotated* rings across time for all trained networks.

#### Uncued colour decoding

For each RNN model, we trained a set of 6 × 2 = 12 linear discriminant analysis (LDA) classifiers (one for each colour category pair and retrocue location) to perform pairwise colour-category discriminations. We first extracted the population hidden activity vectors at the last timestep of the post-cue delay for all stimuli and labelled the data according to the uncued stimulus on each trial. We then binned the labels into B colour categories, as described before. We trained the LDA classifiers in two-fold cross-validation – that is, for each of the 12 conditions, we split the corresponding data into two subsets. We trained and tested a given LDA classifier on the first and second subset, respectively. Then, we reinitialised it and used the reverse subset order for training and testing. The two test decoding accuracy scores for the classifier were then averaged to give a final score for the condition. The test scores were further averaged across the 12 trained classifiers for each model prior to statistical analysis at the group level.

#### Colour discriminability

Hidden activity data from each memory delay was sorted into 4 conditions, according to the cued location: L1-cued, L2-uncued, L1-uncued and L2-cued. For each memory delay, we first performed PCA to obtain the 3-dimensional coordinates corresponding to the colour category representations for each condition. Colour discriminability index (CDI) was then defined as:

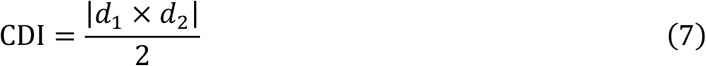

where *d*_1_ and *d*_2_ are vectors corresponding the two diagonals of the data coordinate-defined quadrilateral. The surface area data was averaged across both locations. The pre-cue cued and uncued conditions were also averaged to create a single ‘pre-cue’ category, as the coding scheme was observed to be location- (and not cue-) based at that timepoint. We assessed for differences between the pre-cue, cued and uncued conditions with contrasts based on the results from [10].

#### Learning_dynamics

The plane fitting procedure and AI calculation was identical to that described above. We compared the training loss curves between models and observed that they mostly followed the same shape, characterised by a pronounced plateau period occurring at approximately half the initial loss value (see panel **A** in **S1 Fig**). Therefore, we chose to compare AI values across three training stages: pre-training, at the mid-learning plateau, and after the training had concluded. To find the epoch corresponding to the plateau for each model, we first smoothed the loss curve by applying a Gaussian filter (with standard deviation set to 4) and subsequently calculated its derivative with respect to training time (see panel **B** in **S1 Fig** for an example). The epoch designated as the plateau timepoint was defined as the first local maximum in the derivative curve, constrained to exclude the early epochs where the loss value was within a 5% margin of the initial loss value. This constraint was necessary because all models showed a plateau period at the beginning of the training before the loss started to decrease exponentially.

#### Regression of representational geometry versus training latency

We trained N=100 models using the same hyperparameter settings as described above (in the section *Task structure and training procedures*) but using a different training stop procedure. To equate performance across the trained networks, we stopped training after the loss value averaged over the last epoch fell below the threshold of 0.0005 and noted the total number of training epochs needed to reach that timepoint by each network (time to convergence). Models were then evaluated on the test dataset. We calculated the subspace AI values for the three geometries of interest (*Cued, Uncued, Cued/Uncued*) using the hidden activity patterns from the last timepoint of the post-cue delay. The AI for the *Cued* geometry was computed by substituting **X**^*full*^ entries corresponding to the cued-L1 and cued-L2 activity patterns for S_1_ and S_2_. The AI for the *Uncued* geometry was calculated using an analogous matrix where the data had been sorted and binned by the uncued stimulus colour. Finally, to compute the AI for the *Cued/Uncued* geometry, we first split the test dataset into two subsets according to the location of the retrocue. For each subset, we sorted and binned the trials into B = 4 bins by the cued colour, and then repeated this step using the uncued colour labels. We then averaged within these bins to calculate the mean hidden activity patterns for the cued and uncued colours, which served as S_1_ and S_2_ for the AI analysis. The two AI scores (corresponding to the cued-L1/uncued-L2 and cued-L2/uncued-L1 subspaces) were averaged to obtain a final *Cued/Uncued* score for each network. The *Cued*, *Uncued* and *Cued/Uncued* AI values were entered as the predictor variables in a multiple linear regression model, predicting the time to convergence. The dependent variable was log-transformed prior to being used in the regression model.

### Experiment 2

#### Task structure and training procedures

The overall task structure was the same as for Experiment 1, but we manipulated the lengths of the two delay intervals during training and test. The previously used training set was further expanded to include all combinations of delay lengths (2, 6, 7 and 9 cycles for both pre- and post-cue delay, drawn independently), resulting in a total of 8192 unique trials. The models (N = 30) were not subject to any noise (σ = 0), to investigate the potential effects of the variable delay length manipulation on the learnt task solutions in isolation. We trained the individual networks until the mean loss value from the last epoch crossed the threshold of 0.0005.

#### Behavioural analysis

We created 3 datasets, which differed in the length of the delay interval used. These corresponded to a delay length experienced during training and directly comparable with the first set of trained networks (7 cycles), as well as two novel delay lengths, chosen from within and outside the training range (4 and 10 cycles, respectively).

#### Cross-temporal decoding

For each trial timepoint, we trained a set of 12 LDA classifiers to perform pairwise discriminations between colour categories using the population hidden activity vectors. Data was labelled according to the cued stimulus (labels corresponded to 4 colour categories). Each set of classifiers was subsequently tested on an unseen subset of the data which included all trial timepoints. To assess the stability of memory maintenance dynamics for a given condition (fixed or variable delay), we computed the average off-diagonal delay cross-temporal decoding accuracy scores for all models and tested them against the chance decoding level (50%) with a one-sample t-test. To compare the stability of the delay code between models trained with fixed and variable delay lengths, we calculated the ratio between the off- and on-diagonal cross-temporal decoding accuracy scores from the two delays and compared this metric between the groups with an independent samples t-test.

### Experiment 3

We modified the previously used trial structure by setting the length of the pre- and post-cue delay periods to 5 cycles, as well as including a location probe (1 cycle), and a third, final memory delay of 5 cycles (post-probe delay). We designed 3 experimental conditions, defined by the proportion of trials on which the retro-cue and probe inputs matched: 100%, 75% and 50%. As in Experiment 2, the models (N=30 per condition) were not subject to any noise, and we stopped training once the loss (summed over all the validly cued trials in the last epoch) crossed the threshold of 0.0005. The training dataset included 512 unique trials and for each epoch, we drew a random subset on which the retro-cue and probe inputs were set to mismatch. To increase the statistical power of the multiple linear regression analysis, we increased the sample size to N=100 networks for each group. The Cued, Uncued and Cued/Uncued AI values were calculated by following the same steps as outlined above for Experiment 1 except using the hidden activity data from the last timepoint of the post-probe delay. The dependent variable (total number of training epochs) was box-cox transformed (separately for each validity condition) prior to being used in the multiple regression model.

#### Behavioural analysis

We sampled network choices as previously described and then fit a 3-component mixture model to the trial-wise choice data, separately for every model, using the Analogue Report Toolbox (https://www.paulbays.com/toolbox/) in MATLAB version 2020b (*Mathworks*). The model is fit using maximum likelihood estimation and is described by:

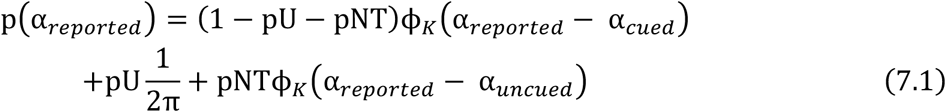

where pU stands for the probability of making random guesses, pNT describes the probability of misremembering the target location and ϕ_*k*_ is a von Mises distribution function with zero mean and concentration parameter *K*. Probability of making the target response (pT) is the difference between 1 and the sum of pU and pNT. Thus, choices are modelled as arising from a mixture of target and non-target-guided components, as well as random guesses. The fit parameters (pT, pU, pNT and K) were compared between trials on which the retro-cue and probe inputs matched (valid) and mismatched (invalid trials), as well as across the experimental conditions (100%, 75% and 50% retro-cue validity). To this effect, for each parameter, we used a 2×3 mixed model ANOVA, with a repeated measures factor of trial type, and a between-subject factor of experimental condition. In cases where the data did not satisfy the assumptions of the mixed model ANOVA, we used a combination of non-parametric tests instead. In such cases, to test the main effect of the repeated measures factor of trial type, we concatenated the data from the two validity conditions and assessed the difference between valid and invalid trials with a Wilcoxon signed-rank test. To test the main effect of the between-subject factor of validity condition, we averaged the data between the valid and invalid trials and compared it between the two validity conditions with a Mann-Whitney U-test. Lastly, to test for an interaction between the trial type and validity condition factors, we calculated the difference between the valid and invalid trials and compared it across the validity conditions with a Mann-Whitney U-test.

#### Software

All statistical analyses described were implemented in python using the scipy.stats package (version 1.7.1), with the exception of the ANOVA, multiple linear regression and Bayesian statistics analyses, which were performed in JASP (version 0.12.2). All networks were trained using the pytorch library (version 1.10.2).

## Statistics abbreviations

*M*: mean
*SEM*: standard error of the mean
*CI*: confidence interval

## Acknowledgements

We wish to thank Michał Wójcik, Dante Wasmuht, Jake Stroud, Andrew Saxe and Paul Muhle-Karbe for useful discussions and comments on the manuscript.

## Supporting Information

S1 Fig. Training dynamics of the networks from Experiment 1.

S2 Fig. Networks trained with variable delay interval lengths show improved temporal generalisation on the task but similar representational geometry as those trained with fixed delay lengths.

S3 Fig. Results of the regression analyses predicting the number of training epochs from AI *Cued*, AI *Cued/Uncued* and AI *Uncued* in Experiment 3.

S4 Note. Results of the ANOVA analyses of mixture model parameters fit to choice data from Experiment 3.

## References

1. Fuster JM. The Prefrontal Cortex—An Update. Neuron. 2001;30: 319–333. doi:10.1016/S0896-6273(01)00285-9

2. Goldman-Rakic PS. Cellular basis of working memory. Neuron. 1995;14: 477–485. doi:10.1016/0896-6273(95)90304-6

3. Durstewitz D, Seamans JK, Sejnowski TJ. Neurocomputational models of working memory. Nat Neurosci. 2000;3: 1184–1191. doi:10.1038/81460

4. Wang X-J. 50 years of mnemonic persistent activity: quo vadis? Trends in Neurosciences. 2021;44: 888–902. doi:10.1016/j.tins.2021.09.001

5. Masse NY, Yang GR, Song HF, Wang XJ, Freedman DJ. Circuit mechanisms for the maintenance and manipulation of information in working memory. Nat Neurosci. 2019;22: 1159–1167. doi:10.1038/s41593-019-0414-3

6. Orhan AE, Ma WJ. Efficient probabilistic inference in generic neural networks trained with non-probabilistic feedback. Nat Commun. 2017;8: 138. doi:10.1038/s41467-017-00181-8

7. Griffin IC, Nobre AC. Orienting Attention to Locations in Internal Representations. Journal of Cognitive Neuroscience. 2003;15: 1176–1194. doi:10.1162/089892903322598139

8. Schmidt BK, Vogel EK, Woodman GF, Luck SJ. Voluntary and automatic attentional control of visual working memory. Perception & Psychophysics. 2002;64: 754–763. doi:10.3758/BF03194742

9. Myers NE, Stokes MG, Nobre AC. Prioritizing Information during Working Memory: Beyond Sustained Internal Attention. Trends in Cognitive Sciences. 2017;21: 449–461. doi:10.1016/j.tics.2017.03.010

10. Panichello MF, Buschman TJ. Shared mechanisms underlie the control of working memory and attention. Nature. 2021;592: 601–605. doi:10.1038/s41586-021-03390-w

11. Hajonides JE, Nobre AC, van Ede F, Stokes MG. Decoding visual colour from scalp electroencephalography measurements. NeuroImage. 2021;237: 118030. doi:10.1016/j.neuroimage.2021.118030

12. Flesch T, Juechems K, Dumbalska T, Saxe A, Summerfield C. Rich and lazy learning of task representations in brains and neural networks. bioRxiv. 2021.

13. Libby A, Buschman TJ. Rotational dynamics reduce interference between sensory and memory representations. Nat Neurosci. 2021;24: 715–726. doi:10.1038/s41593-021-00821-9

14. Nieh EH, Schottdorf M, Freeman NW, Low RJ, Lewallen S, Koay SA, et al. Geometry of abstract learned knowledge in the hippocampus. Nature. 2021; 1–5. doi:10.1038/s41586-021-03652-7

15. Xie Y, Hu P, Li J, Chen J, Song W, Wang X-J, et al. Geometry of sequence working memory in macaque prefrontal cortex. Science. 2022;375: 632–639. doi:10.1126/science.abm0204

16. Bernardi S, Benna MK, Rigotti M, Munuera J, Fusi S, Salzman CD. The Geometry of Abstraction in the Hippocampus and Prefrontal Cortex. Cell. 2020; S0092867420312289. doi:10.1016/j.cell.2020.09.031

17. Sheahan H, Luyckx F, Nelli S, Teupe C, Summerfield C. Neural state space alignment for magnitude generalization in humans and recurrent networks. Neuron. 2021;109: 1214–1226.e8. doi:10.1016/j.neuron.2021.02.004

18. Luyckx F, Nili H, Spitzer B, Summerfield C. Neural structure mapping in human probabilistic reward learning. Elife. 2019;8. doi:10.7554/eLife.42816

19. Remington ED, Narain D, Hosseini EA, Jazayeri M. Flexible Sensorimotor Computations through Rapid Reconfiguration of Cortical Dynamics. Neuron. 2018;98: 1005–1019.

20. Elsayed GF, Lara AH, Kaufman MT, Churchland MM, Cunningham JP. Reorganization between preparatory and movement population responses in motor cortex. Nature Communications. 2016;7: 13239. doi:10.1038/ncomms13239

21. Ester EF, Nouri A, Rodriguez L. Retrospective Cues Mitigate Information Loss in Human Cortex during Working Memory Storage. J Neurosci. 2018;38: 8538–8548. doi:10.1523/JNEUROSCI.1566-18.2018

22. Sprague TC, Ester EF, Serences JT. Restoring Latent Visual Working Memory Representations in Human Cortex. Neuron. 2016;91: 694–707. doi:10.1016/j.neuron.2016.07.006

23. Russin J, Zolfaghar M, Park SA, Boorman E, O’Reilly RC. A Neural Network Model of Continual Learning with Cognitive Control. arXiv:220204773 [cs, q-bio]. 2022 [cited 7 Mar 2022]. Available: http://arxiv.org/abs/2202.04773

24. Orhan AE, Ma WJ. A diverse range of factors affect the nature of neural representations underlying short-term memory. Nat Neurosci. 2019;22: 275–283. doi:10.1038/s41593-018-0314-y

25. Bays PM, Catalao RFG, Husain M. The precision of visual working memory is set by allocation of a shared resource. J Vis. 2009;9: 7.1–11. doi:10.1167/9.10.7

26. Flesch T, Juechems K, Dumbalska T, Saxe A, Summerfield C. Orthogonal representations for robust context-dependent task performance in brains and neural networks. Neuron. 2022;110: 1258–1270.e11. doi:10.1016/j.neuron.2022.01.005

27. Flesch T, Balaguer J, Dekker R, Nili H, Summerfield C. Comparing continual task learning in minds and machines. Proceedings of the National Academy of Sciences. 2018;115: E10313–E10322. doi:10.1073/pnas.1800755115

28. Souza AS, Oberauer K. In search of the focus of attention in working memory: 13 years of the retro-cue effect. Atten Percept Psychophys. 2016;78: 1839–1860. doi:10.3758/s13414-016-1108-5

29. Murray AM, Nobre AC, Clark IA, Cravo AM, Stokes MG. Attention restores discrete items to visual short-term memory. Psychol Sci. 2013;24: 550–556. doi:10.1177/0956797612457782

30. Williams M, Hong SW, Kang M-S, Carlisle NB, Woodman GF. The benefit of forgetting. Psychonomic Bulletin & Review. 2013;20: 348–355. doi:10.3758/s13423-012-0354-3

31. Souza AS, Rerko L, Lin H-Y, Oberauer K. Focused attention improves working memory: implications for flexible-resource and discrete-capacity models. Atten Percept Psychophys. 2014;76: 2080–2102. doi:10.3758/s13414-014-0687-2

32. Souza AS, Rerko L, Oberauer K. Getting more from visual working memory: Retro-cues enhance retrieval and protect from visual interference. Journal of Experimental Psychology: Human Perception and Performance. 2016;42: 890–910. doi:10.1037/xhp0000192

33. Gunseli E, van Moorselaar D, Meeter M, Olivers CNL. The reliability of retro-cues determines the fate of noncued visual working memory representations. Psychon Bull Rev. 2015;22: 1334–1341. doi:10.3758/s13423-014-0796-x

34. Makovski T, Pertzov Y. Attention and memory protection: Interactions between retrospective attention cueing and interference. The Quarterly Journal of Experimental Psychology. 2015;68: 1735–1743. doi:10.1080/17470218.2015.1049623

35. van Moorselaar D, Gunseli E, Theeuwes J, Olivers CNL. The time course of protecting a visual memory representation from perceptual interference. Frontiers in Human Neuroscience. 2015;8. doi:10.3389/fnhum.2014.01053

36. Wallis G, Stokes M, Cousijn H, Woolrich M, Nobre AC. Frontoparietal and Cingulo-opercular Networks Play Dissociable Roles in Control of Working Memory. J Cogn Neurosci. 2015;27: 2019–2034. doi:10.1162/jocn_a_00838

37. Myers NE, Stokes MG, Nobre AC. Prioritizing Information during Working Memory: Beyond Sustained Internal Attention. Trends Cogn Sci (Regul Ed). 2017;21: 449–461. doi:10.1016/j.tics.2017.03.010

38. Saxe AM, McClelland JL, Ganguli S. Exact solutions to the nonlinear dynamics of learning in deep linear neural networks. arXiv; 2014 Feb. Report No.: arXiv:1312.6120. doi:10.48550/arXiv.1312.6120

