## Supplementary material for "A recurrent neural network model of prefrontal brain activity during a working memory task": S1

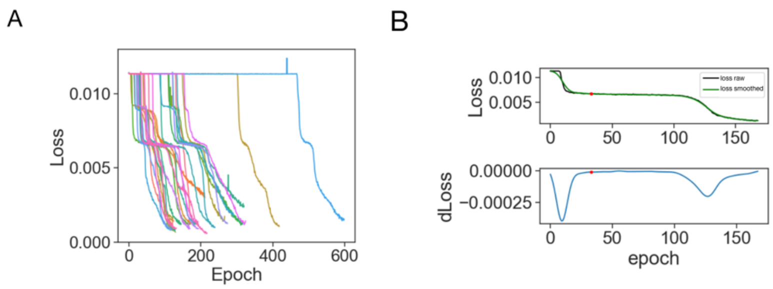


**S1 Fig. Training dynamics of the networks from Experiment 1. A.** Training loss values across all epochs, colours correspond to individual networks. **B.** *Upper:* Raw (black) and smoothed (green) training loss plotted across epochs. *Lower:* The derivative of the smoothed loss with respect to time. Red dot in both panels corresponds to the local minimum chosen as the ‘mid-training plateau’ timepoint (refer to the *Methods* section in the main text for details). Data from an example model.
