## Supplementary material for "A recurrent neural network model of prefrontal brain activity during a working memory task": S2

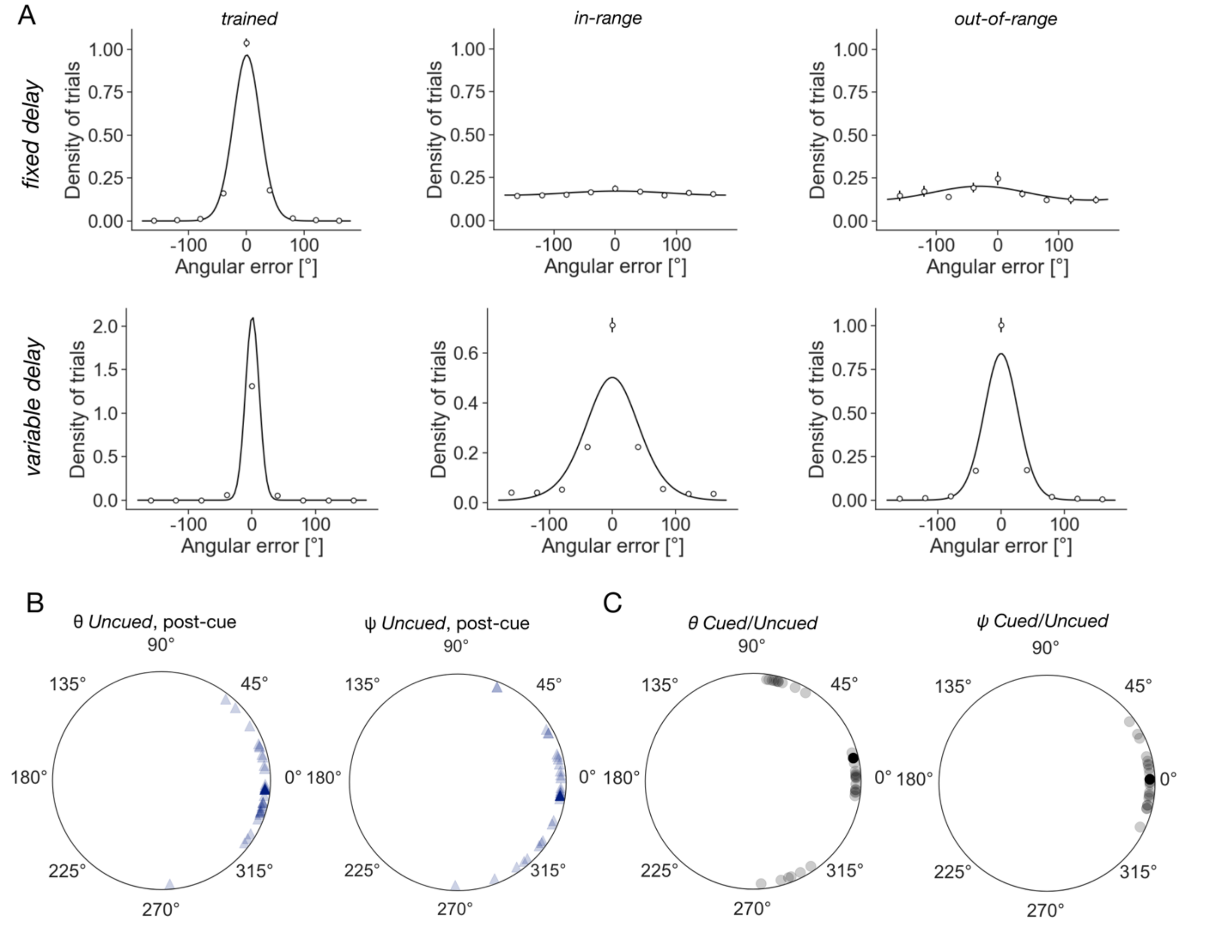


**S2 Fig. Networks trained with variable delay interval lengths show improved temporal generalisation on the task but similar representational geometry as those trained with fixed delay lengths. A.** Angular error density plots for the 3 different test conditions: datasets with the delay interval length used during training (*trained*), a previously not experienced delay length that fell within the training range (*in-range*), and a novel delay outside of the training range (*out-of-range*). First and second rows correspond to the data from the networks trained with a fixed delay and variable delay intervals, respectively. Datapoints correspond to *M*±*SEM* across all models, shown with the best von Mises fit (solid line). **B.-C.** Angles $\theta$ and ψ between the two *Uncued* and *Cued*/*Uncued* planes (averaged across the two retrocue locations) in the post-cue delay interval, respectively. Individual models shown in transparent, grand averages in opaque colours.
