## Supplementary material for "A recurrent neural network model of prefrontal brain activity during a working memory task": S3

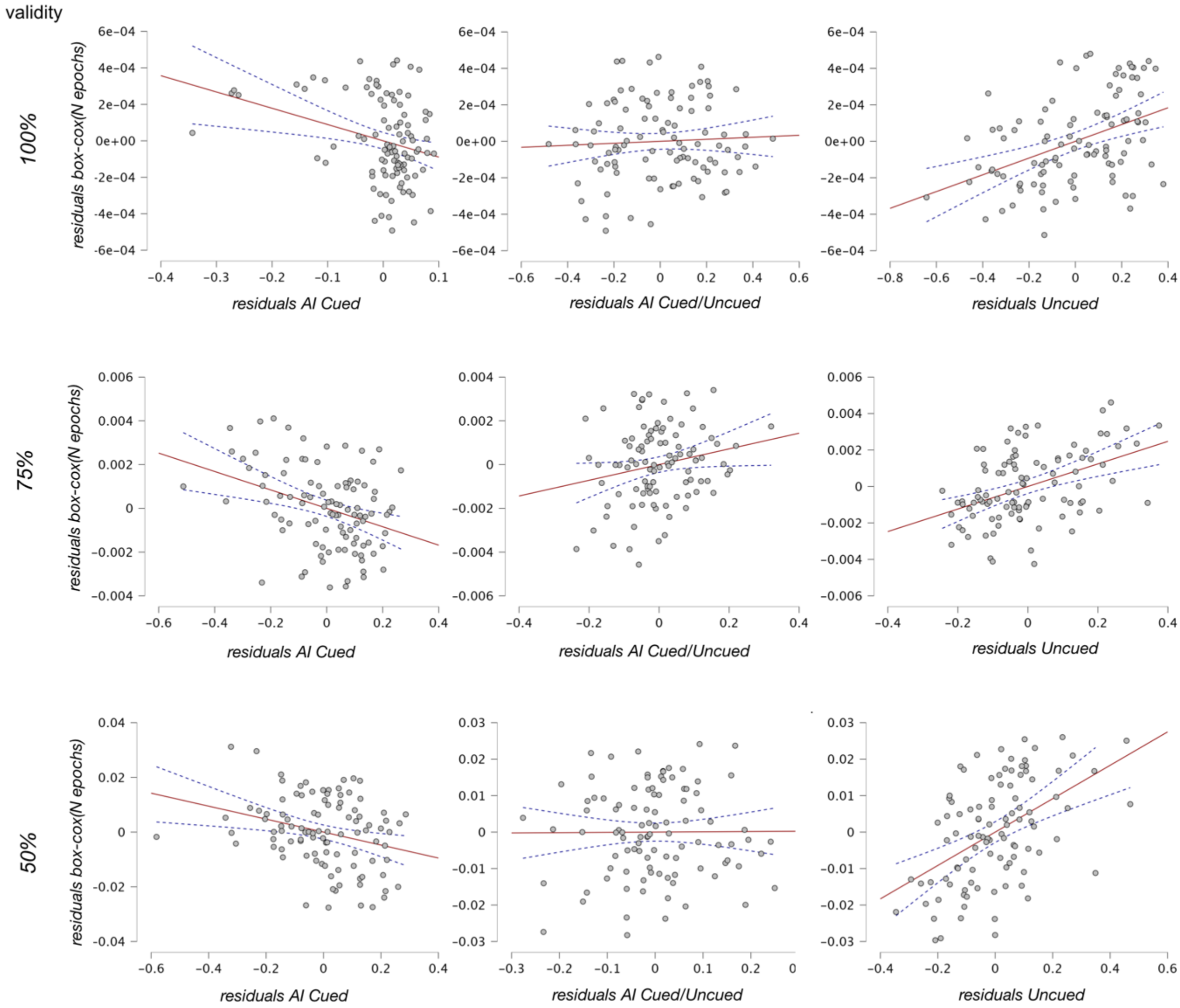


**S3 Fig.** Results of the regression analyses predicting the number of training epochs (after a box-cox transformation) from AI *Cued*, AI *Cued/Uncued* and AI *Uncued* (columns, left to right*)* in Experiment 3. Rows correspond to the three retrocue validity conditions examined (100, 75 and 50%, top to bottom). Data shown in grey dots, regression lines of best fit in red, 95% confidence interval for the slope in navy dashed lines.
