## Supplementary material for "A recurrent neural network model of prefrontal brain activity during a working memory task": S4

**S4 Note. Results of the ANOVA analyses of mixture model parameters fit to choice data from Experiment 3.**

As noted in the main text, we compared the mixture model parameters fit to the choice data generated by models from Experiment 3 with 2x2 mixed model ANOVAs (or non-parametric equivalents in the case of the probability parameters). The full results of this analysis are described below (data shown in **Fig. 6C).**

The memory precision parameter (*K*) showed a significant main effect of trial type ($F(1,58) = 144.91$, $p < .001$; valid = 26.9, invalid = 22.78) and a significant interaction with condition ($F(1,58) = 144.76$, $p < .001$), but no main effect of condition ($F(1,58) = 0.86$, $p = .358$). Post-hoc tests with Holm-correction revealed that recall precision was significantly higher on valid than invalid trials in the 75% validity condition ($t(29) = 17.02$, $p < .001$), whilst there was no difference in the 50% validity condition ($t(29) = .004$, $p = .997$). Furthermore, the precision parameter on invalid trials was significantly lower in the former compared to latter condition ($t(29) = -4.22$, $p < .001$), with an opposite trend on valid trials, which did not however reach statistical significance ($t(29) = 2.40$, $p = .057$).

An analogous analysis concerning the probability of recalling the target item (*pT*) revealed that it was significantly higher on valid than invalid trials (Wilcoxon signed-rank test, $W(59) = 1337$, p = .002; valid = .996, invalid = .994) as well as in the 75%, compared to 50% validity condition (Mann-Whitney U-test: $W(29) = 286$, $p = .015$; $pT = .997$ and $pT = .992$, respectively). There was also a significant interaction between the two factors (mean difference between valid and invalid trials – 75% validity: .003, 50% validity: $6.02 \times 10^{-6}$, Mann-Whitney U-test: $W(29) = 208$, $p < .001$). Holm-corrected post-hoc tests revealed that *pT* was significantly higher on valid trials in the probabilistic than in the neutral condition (Mann-

Whitney U-test: $W(58) = 199$, $p < .001$), with no significant difference between the conditions on invalid trials (Mann-Whitney U-test: $W(58) = 336$, $p = .187$). In the 50% validity condition, there was no significant difference between the two trial types (Wilcoxon signed-rank test: $W(29) = 224$, $p = .871$). In contrast, in the 75% validity condition, *pT* was significantly higher on valid than invalid trials (Wilcoxon signed-rank test: $W(29) = 430$, $p < .001$).

Probability of random guesses (*pU*) was significantly lower on valid than invalid trials (Wilcoxon signed-rank test, $W(59) = 571$, $p = .011$) and in the 75%, compared to the 50% validity condition (Mann-Whitney U-test: $W(29) = 618$, $p = .013$). The interaction between the trial type and condition was also significant, with a more pronounced difference between valid and invalid trials in the 75% validity condition (Mann-Whitney U-test: $W(29) = 637$, $p = .005$; $pU = -.002$ and $pU = -7.810 \times 10^{-5}$ for 75% and 50% validity, respectively). Holm-corrected post-hoc tests revealed a significant between-condition difference on valid (Mann-Whitney U-test: $W(58) = 683$, $p = .002$, $pU = 0.007$ and 0.001 for 50 and 75% validity, respectively) but not invalid trials (Mann-Whitney U-test: $W(58) = 588$, $p = .084$). In the 75% validity condition, *pU* was significantly lower on valid than invalid trials (Wilcoxon signed-rank test, $W(29) = 75$, $p = .004$; $pU = 0.001$ and 0.003, respectively), whilst there was no such difference in the 50% validity condition (Wilcoxon signed-rank test, $W(29) = 234$, $p = .984$).

Lastly, the analogous analysis regarding the probability of recalling the non-target item (*pNT*) yielded a significant main effect of trial type (Wilcoxon signed-rank test, $W(59) = 577$, $p = .013$, valid = $4.68 \times 10^{-4}$, invalid = $8.89 \times 10^{-4}$) but no main effect of the validity condition (Mann-Whitney U-test: $W(29) = 447$, $p = .971$). The interaction between the two factors was significant (mean difference in *pNT* between valid and invalid trials – 75% validity: $-9.14 \times 10^{-4}$, 50% validity: $7.0 \times 10^{-5}$, Mann-Whitney U-test: $W(29) = 628$, $p = .008$). Holm-corrected post-hoc tests revealed a significant between-condition difference on valid (Mann-Whitney U-test: $W(58) = 659$, $p = .006$, $pNT = 6.65 \times 10^{-4}$ and 0.001 for 50 and 75% validity, respectively), but not invalid trials (Mann-Whitney U-test: $W(58) = 410$, $p = .562$). In the 75% validity condition, *pNT* was significantly lower on valid than invalid trials (Wilcoxon signed-rank test, $W(29) = 75$, $p = .002$; $pNT = 1.99 \times 10^{-4}$ and .001, respectively), whilst there was no such difference in the 50% validity condition (Wilcoxon signed-rank test, $W(29) = 225$, $p = .887$).

In summary, we found significant interaction effects between the validity condition and trial type for all mixture model parameters examined. In the 50% validity condition, there were no significant trial-type differences in the fitted parameters. In contrast, in the 75% validity condition, both the precision (*K*) and probability of recall (*pT*) of target items were significantly higher on valid than invalid trials, whilst the probability of recalling the non-target item (*pNT*) and making random guesses (*pU*) followed the opposite pattern. It is however worth noting that all the estimated probability quantities were close to ceiling (or floor, in the case of *pNT* and *pU*).
